# Commensal fungi *Candida albicans* modulates dietary high-fat induced alterations in metabolism, immunity, and gut microbiota

**DOI:** 10.1101/2022.03.23.485455

**Authors:** Doureradjou Peroumal, Satya Ranjan Sahu, Premlata Kumari, Bhabasha Utkalaja, Narottam Acharya

**Author notes:** Correspondence to: Narottam Acharya, Phone: -674-230 0728. **Abbreviations:** HFD: high-fat diet; ND: Normal diet; GLP-1: glucagon-like peptide-1; GIP: Glucose- dependent insulinotropic polypeptide; PYY: peptide tyrosine tyrosine; PP: pancreatic polypeptide; DIO: Diet induced obesity; RBG: Random blood glucose.

## Abstract

Obesity and its associated diseases represent a growing health concern. Despite a strong correlation between gut microbiota and obesity, a limited study is available to suggest the direct involvement of fungus in host physiology. *Candida albicans* survives as a commensal fungus in the gastrointestinal tract, and its excessive growth causing infections in immune-suppressed individuals is widely accepted. However, any mutualistic relationship that may exist between *C. albicans* and the host remains outstanding. Here we show that the dietary *C. albicans* do not cause any noticeable infections, and our metagenomics analyses suggest that it colonizes in the gut and modulates microbiome dynamics which in turn negates the high-fat diet induced uncontrolled body weight gain, metabolic hormonal imbalances, and inflammatory response. Interestingly, adding *C. albicans* to a non-obesogenic diet stimulates the appetite regulated hormones and helped the mice gain healthy body weight. In concert, our results suggest a mutualism between *C. albicans* and the host, therefore, contrary to the notion, *C. albicans* is not always an adversary rather a bonafide admirable companion of the host. Finally, we discuss its potential translational implication as a probiotic especially in obese people or people dependent on high fat calorie intakes to manage obesity and associated complications.

## Introduction

In 2019, World Health Organization reported that about 650 million adults and 38 million children under the age of five were overweight or obese worldwide. Moreover, an inactive lifestyle of people due to unexpected lockdowns by the current COVID19 pandemic and their dependence on a high-fat and sugary diet might have substantially increased the present obese populations. The mortality rates in overweight and obese subjects are very high than the underweight subjects. Thus, obesity is a major social public health concern worldwide given its link to chronic diseases like diabetes, cancer, cardiovascular, and neurological disorders associated with fatality[1, 2]. In addition, influenza, bronchitis, pneumonia, tuberculosis, and other nosocomial infections also contribute substantially to morbidity and mortality in obese individuals. High-fat calorie consumption deregulates polyunsaturated fatty acids compositions, which are the proven precursors for several mediators of chronic inflammation[3]. An increase in adipose tissue (AT) mass is a dominant attribute of obesity. While AT-resident immune cells play important roles in maintaining AT homeostasis, obesity alters their numbers and activities, which modulate inflammatory responses[4]. Reduction in anti-inflammatory immune cells and the accumulation of pro-inflammatory immune cells in AT are crucial in the pathogenesis of obesity and its metabolic complications[5]. Interestingly, in experimental models as well as human obese subjects, an accumulation of these activated immune cells in the circulating pool and other tissues including the liver, hypothalamus, lungs, skeletal muscles, etc. were also noticed[6–8].

A large body of evidence suggested that gut microbiota composed of bacteria, archaea, virus, and fungi as the main biotic factor regulate obesity, metabolic syndrome, type 2 diabetes, immunological, and neurological status of its human host[9–12]. In addition to maternal transmission during birth, food ingestion is another way microorganisms enter the gastrointestinal tract and diet is one of the factors that influence microbial diversity and richness[13]. The gut microbiota of healthy individuals is mostly composed of bacteria from Firmicutes and Bacteroidetes. The proportion of these bacteria often gets altered in obese animals and humans. Recent studies also indicate the role of the fungal population, also known as mycobiota, in nutrient metabolism and the intestinal health of humans and animals. Human gut mycobiome harbors more than 66 genera and 184 species of fungi, while *Candida*, *Saccharomyces,* and *Cladosporium* predominate. Although no differences were detected in the richness of the mycobiome between non-obese and obese subjects, family biodiversity was significantly reduced in obese subjects compared to that in non- obese individuals[14]. *Candida* sp. survive mostly as commensals in the intestine of healthy individuals. Although some studies have suggested a link between expansion in *Candida* sp. and diabetes and inflammatory disorders of the gastrointestinal tract with a possible active role for *Candida albicans*, the difference in the relative abundance of *C. albicans* in obesity was not observed[15, 16]. Contrarily, a recent study points out an association of *Candida albicans, Candida kefyr, and Rhodotorula mucilaginosa* fungal species with human obesity[17].

Current approaches for the treatment of obesity, including diet and lifestyle changes, have not been successful. Improved understanding of microbiome including mycobiome, and their association with obesity may yield new treatment measures for obesity-associated disorders. The transplantation of microbiome isolated from fecal samples of non-obese mice controlling metabolic syndromes in obese mice implements the role of these microorganisms in obesity and has given a new potential dimension to manage obesity[18]. Meanwhile, as some of the microorganisms are non-culturable, the application of the microbiome to control obesity seems not to be a viable option. The number of uncultured bacterial species in the gut microbiota currently stands at 1952[19]. Therefore, a successful experimental demonstration of the role of these fungal or bacterial species individually or in a mixture in regulating obesity will be a comprehensive approach with better translational potential. On that line, since *Candida albicans* is a predominant *Candida* species and their association with obesity in the context of mycobiome has been appreciated, we wanted to decipher its role in the regulation of host’s obesity, immunity, and microbiome in an animal model. Secondly, *C. albicans* has been explored mostly as a fungal pathogen to cause superficial and systemic candidiasis and being a commensal, its beneficial attributes to the host remain unexplored. Our findings suggest that supplementation of *C. albicans* to the food of mice antagonizes the deleterious effects of a high-fat diet and restores the alterations in metabolism and immunity induced by the obesogenic diet by changing the gut microbiota to a healthy standard.

## Results

### Colonization of dietary *C. albicans* in the gut does not induce body weight gain while it protects mice from high-fat diet induced obesity

We preferred BALB/c mice, despite different obesity models[20–23] as it is assumed to be obesity resistant and highly susceptible to *C. albicans* systemic infection ^26,28,29,^ an ideal system to determine the effect of dietary form of *C. albicans* on host physiology, obesity and if any infections caused thereafter. Thus, the study involved similar housed four groups of male mice matched for age, bodyweight, and random blood glucose (n=8 per group on average) were subjected to normal diet (ND) or obesogenic high-fat diet (HFD) mixed with or without *C. albicans* over a period of 150 days (**Fig. 1A**). On day 0, the body weight of mice in each group ranged from 23.1 ± 0.5 to 24.4 ± 0.6 grams (**Supp. Table 1**). Up to 90 days of dietary intervention, the percent body weight gain was similar in each group and parallel to the BALB/c diet induced obesity (DIO) model in previous reports, suggesting the use of similar experimental conditions among these studies[20, 24, 25] (**Fig. 1B**). Subsequently, the percent body weight doubled after every eight weeks of HFD consumption alone, with an average body weight of ∼36 gms on the 150th day, an increase of ∼50% compared to a normal chow-fed group that gained only ∼25% for the period over their initial weight (**Fig. 1B and Suppl. Fig.1A**). However, the presence of *C. albicans* culture in the diets had an opposing influence on body weight gain of normal and HFD fed mice. Interestingly, while the presence of *C. albicans* improved the normal chow-fed mice in gaining weight marginally, it considerably reduced the excessive body weight gain in HFD fed mice. Only an increase of ∼35% body weight gain was found in mice subjected to *C. albicans* mixed HFD as opposed to ∼50% increase in HFD alone fed mice. In addition to an increase in body weight, the presence of visceral and epididymal fat that implicates adiposity was observed maximally in the abdomen of HFD fed mice compared to other groups (**Fig. 1C**). *C. albicans* mixed HFD group displayed relatively less visceral fat depositions. Though the ND fed mice preserved a negligible amount of abdominal fat, interestingly *C. albicans* mixed ND group had relatively a higher deposition of epididymal fat without contributing to significant weight gain (**Fig. 1C**). Generally, gut and feces share similar microbial populations [26]. To understand these effects were due to gut colonization of *C. albicans*, fecal metagenomic DNA was isolated and *C. albicans* presence was confirmed by PCR using specific primers for full length CaPCNA (DNA clamp in replication) and a partial fragment of CaEfg1 (a transcription factor involved in filamentation) orfs [27–29]. Interestingly, both the PCR products were detected in the meta-DNA of mice subjected to a normal diet suggesting that the mice gut is a natural habitat of *C. albicans*, albeit with less abundance (lane 4). However, a substantial amount of PCR products were detected in other samples and the maximum amount was detected in HFD fed mice (compare lanes 3 with 1, 2, and 4) (**Fig. 1D**) indicating counteractive effects of dietary compositions and *C. albicans* gut colonization. Further, the presence of *C. albicans* in the gut was confirmed by estimating the colony forming units (CFUs) efficiency of various fecal samples (data not shown). Thus, these results signify to a possible beneficial effect of *C. albicans* on host metabolic status impacting obesity and metabolic syndrome depending on the dietary fat level and warrant a detailed investigation.

**Figure 1.**
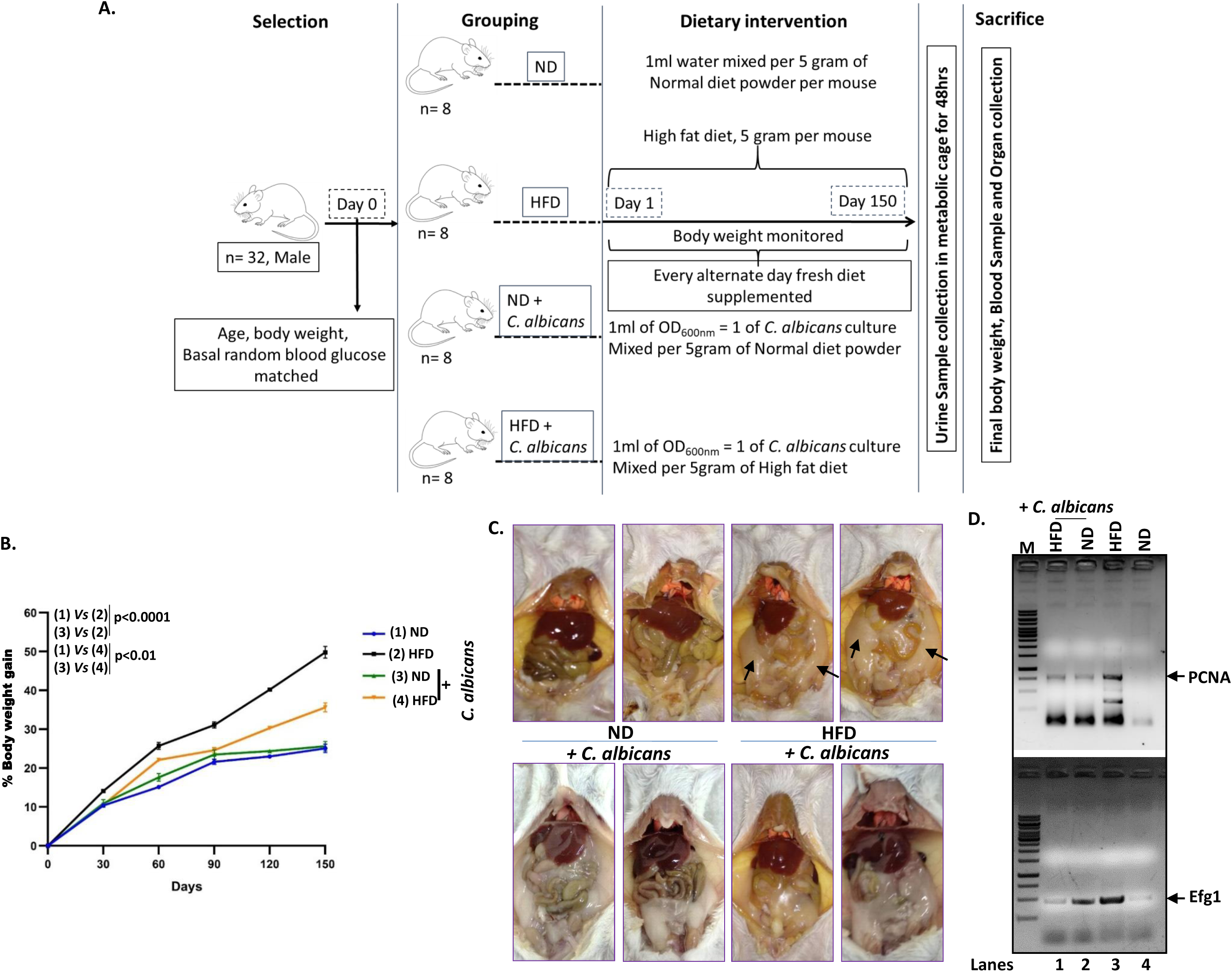
DOI induced body weight gain, fat deposits and fungal colonization in male BLAB/c mice. (**A**) A schematic workflow chart of the study of diet induced obesity and the role of *C. albicans* on weight gain in mice. (**B**) A kinetics of percent gain in body weight with respect to duration (corrected from basal body weight in grams) from mice fed with or without *C. albicans* in normal diet (ND) or high-fat diet (HFD), a statistical significance (*p≤0.05, ** p≤0.01,***p≤0.001, **** p≤0.0001) was calculated using two-way ANOVA and Tukey’s multiple comparisons test. (**C**) A representative couple of images of mesenteric fat indicated by arrows from each groups of mice under dissection are displayed (4 images above are from respective diet without *C. albicans* while 4 images below are with respective diet with *C. albicans*). (**D**) PCR amplification of CaPCNA and CaEfg1 orfs from the meta-genomic DNA isolated from fecal samples. Lanes 1, DNA ladder, 2, HFD + *C. albicans*; 3, ND + *C. albicans*; 4, HFD; and 5, ND.

### Dietary *C. albicans* reduced adiposity associated hormones in diet induced obesity mice

Leptin and resistin play important role in obesity and type II diabetes mellitus (T2DM), together with gastrointestinal ghrelin regulate energy balance and body weight[30, 31]. DIO and genetic models of obesity reported increased leptin and resistin levels. Our estimation revealed that plasma leptin (14-20 ng/ml) and resistin (20-32ng/ml) levels in HFD fed mice were ∼7-10 folds and ∼1.8 - 2.5 folds higher than their respective levels in the normal diet fed mice (1-3 ng/ml) and (7.7-17.2 ng/ml) (**Fig. 2A and B**). Interestingly, *C. albicans* in HFD significantly reduced both leptin 4-6 folds (2.5-5.5 ng/ml) and resistin 1.5 – 1.7 (13.9 - 18.5 ng/ml) than the HFD fed alone. However, leptin and resistin levels were found similar in mice subjected to a diet mixed with fungus regardless of calorie content, suggesting a direct or indirect role of dietary *C. albicans* in controlling the hormones. Further a linear regression analysis revealed a strong positive correlation of body weight gain (grams) versus (ng/ml) leptin (r=0.749; p≤ 0.0001) or resistin (r=0.534; p≤ 0.007) level in each mouse irrespective of treatment background (**Fig. 2D and E**). Ghrelin hormone stimulates appetite, increases food intake, and promotes fat storage in the reproductive system. Acylated ghrelin (active ghrelin) in plasma of ND (16.5 -68.1 pg/ml), HFD (74.5 -153.5 pg/ml), and *C. albicans* mixed HFD (74.57 – 169 pg/ml) fed mice groups were not statistically differed but markedly increased by ∼16 folds in *C. albicans* mixed ND (355 -544 pg/ml) group of mice (**Fig. 2C**). Thus increased active ghrelin level, along with epididymal fat deposition without changing body weight (**Fig. 1C**) in *C. albicans* mixed ND fed mice indicated the possible role of dietary *C. albicans* on host reproductive capacity, in addition to increase feeding tendency by maintaining hunger sense. Further, estimation of leptin/ghrelin ratio as a predictor of body mass index (BMI) as reported earlier in human studies[32, 33] was found to be significantly high in HFD fed mice (∼7 fold high) than ND fed mice, but it was significantly reduced to ∼3.5 folds by *C. albicans* dietary intervention. *C. albicans* addition to ND did not affect on the leptin/ghrelin ratio (**Suppl. Fig.1B**). These results strongly suggest that HFD alone induced the onset of obesity in BALB/c mice but dietary *C. albicans* inhibited the process.

**Figure 2.**
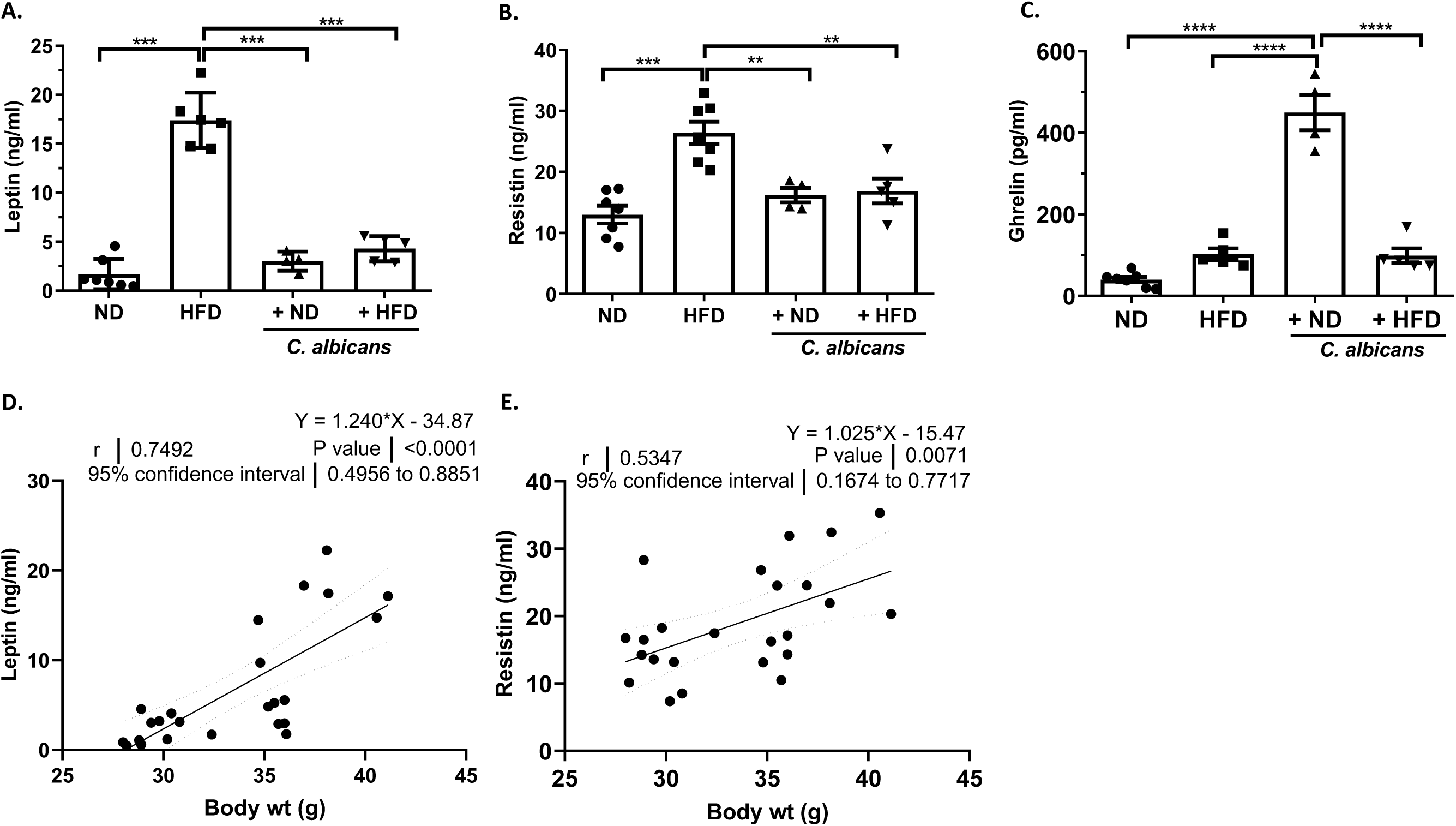
Obesity induced hormonal imbalance associated with body weight. Mean and standard error mean of plasma (**A**) leptin, (**B**) resistin and (**C**) ghrelin level from each of four groups of mice, a statistical significance (*p≤0.05, ** p≤0.01,***p≤0.001, **** p≤0.0001) was calculated using one-way ANOVA and Tukey’s multiple comparisons test. A regression and Person’s correlation analysis of body weight versus (**D**) leptin and (**E**) resistin level from individual mouse irrespective of dietary intervention.

### Altered hormones of energy metabolism in DIO were restored upon feeding *C. albicans* in diet

The roles of gastrointestinal (GI) peptide hormones in the regulation of GI function and energy balance have been demonstrated [34, 35]. In contrast to ghrelin, a low level of glucagon-like peptide 1 (GLP-1) stimulates more eating thereby increasing the probability of obesity onset. GLP-1 was significantly high (0.3 -0.7 ng/ml) in the ND fed mice irrespective of *C. albicans* presence, suggesting them to be low appetite groups. Contrastingly, GLP-1(∼0.1 ng/ml) was significantly low in the HFD group indicating a high tendency to gain more weight (**Fig. 3A**). Like GLP-1, GIP is also an incretin that is secreted in the proximal small intestine in the presence of nutrients in the gut and affects insulin secretion in response to a meal. GIP is also involved in lipid metabolism and is thought to promote fat deposition. GIP levels were elevated in the normal and obesogenic fed mice group (3-4.5 ng/ml) and they were greatly reduced when the feed was blended with *C. albicans* culture (1-2 ng/ ml) (**Fig. 3B**). Pancreatic polypeptide (PP) is a polypeptide secreted by PP cells in the pancreas and its level is reduced in conditions associated with increased food intake but elevated in anorexia nervosa. On fasting, plasmic PP concentration is 80 pg/ml, and after the meal, it raises by 8 to 10 times[36]. The plasma PP levels were significantly reduced in the HFD diet containing *C. albicans* fungal mix (∼40 pg/ml) in comparison to other groups of mice (∼60 pg/ml) (**Fig. 3C**). However, the concentration of Peptide tyrosine tyrosine (PYY), an anorexigenic peptide released from the cells in the ileum and colon, did not alter in any group of mice (∼200-300 pg/ml) (**Suppl. Fig. 1C**). C-peptide, insulin, glucagon, and amylin are released from the pancreas and regulate plasma glucose homeostasis. Insulin and C-peptide levels (∼6 ng/ml and 12 ng/ ml, respectively) were very high in the blood of mice subjected to an obesogenic HFD diet and thus they were appeared to be insulin resistant and hyperglycemic, but in the presence of *C. albicans* mixed food reduced them to respective standard levels (∼2 ng/ml and 5 ng/ ml, respectively) (**Fig. 3 D and E**). Amylin and glucagon level did not alter much in any of the experimental setups used and they remained constant at 100-150 pg/ml of blood **(Suppl. Fig. 1 D and E)**. A direct correlation between leptin and insulin released was also observed (**Suppl. Fig. 1 F and G**). These results suggested that the addition of *C. albicans* to the regular diet increases the hunger hormone level ghrelin but reduces GIP, the appetite inducing hormone; thereby it helps to maintain body weight and prevents overweight. Rising levels of insulin and C-peptide levels in HFD fed mice and their restoration by the presence of dietary *C. albicans* suggested that this commensal fungus is just not a passive component of the gut but it plays an active role in regulating metabolic conditions that predispose to obesity and diabetes.

**Figure 3.**
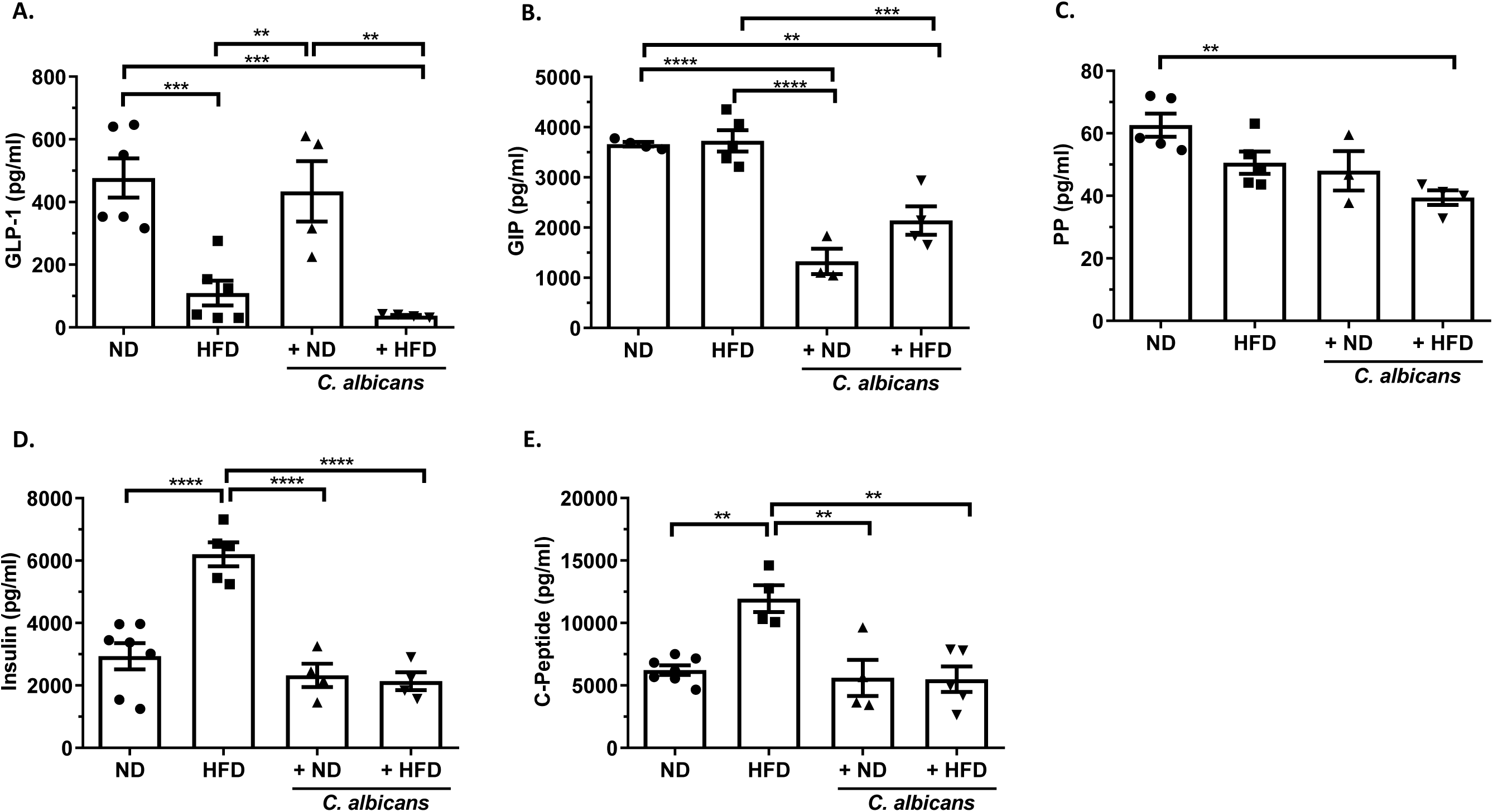
Key metabolic hormonal imbalance associated with energy metabolism and obesity. Mean and standard error mean of various metabolic hormones from each of the groups of mice blood on the 150th day (∼22 weeks) of dietary intervention, a statistical significance (*p≤0.05, ** p≤0.01,***p≤0.001, **** p≤0.0001) was calculated using one-way ANOVA and Tukey’s multiple comparison test. (**A**) glucagon-like peptide-1(GLP-1) level (pg/mL), (**B**) Glucose-dependent insulinotropic polypeptide (GIP) level (pg/mL), (**C**) pancreatic polypeptide (PP) (pg/mL), (**D**) Insulin (pg/mL), and (**E**) C-peptide (pg/mL).

### Dietary *C. albicans* suppresses diet induced alteration in glucose and lipid metabolism

Next, we estimated random blood glucose (RBG) by GlucoOne blood glucose monitoring system and urine sugar by Benedict’s solution reaction to determine any alterations in glucose metabolism in these mice. The RBG levels in the range of 80-140 mg/dl before food and 100-160 mg/dl after taking food are considered to be normal. On the 150^th^ day, RBG levels were highly variable among the animals irrespective of their diet intakes, and it varied from 125 mg/dl to 170 mg/dl **(Fig. 4 A)**. While no traces of glucose was observed in the urine of normal chow-fed mice, in HFD fed mice, Benedict’s solution turned instantly into green by the addition of urine indicating the presence high level of reducing sugars **(Fig. 4 B and Supp. Fig. 2A)**. The bluish-green coloration of urine of mice fed with any diet but mixed with *C. albicans* culture suggested the presence of traces of sugars in their urines. Although we did not find a change in the sugar levels in the blood of various mice groups, excess sugar in the urine of HFD fed mice indirectly indicated a high level of sugars in the blood of those mice, which was reduced by providing *C. albicans* mix to the diet.

**Figure 4.**
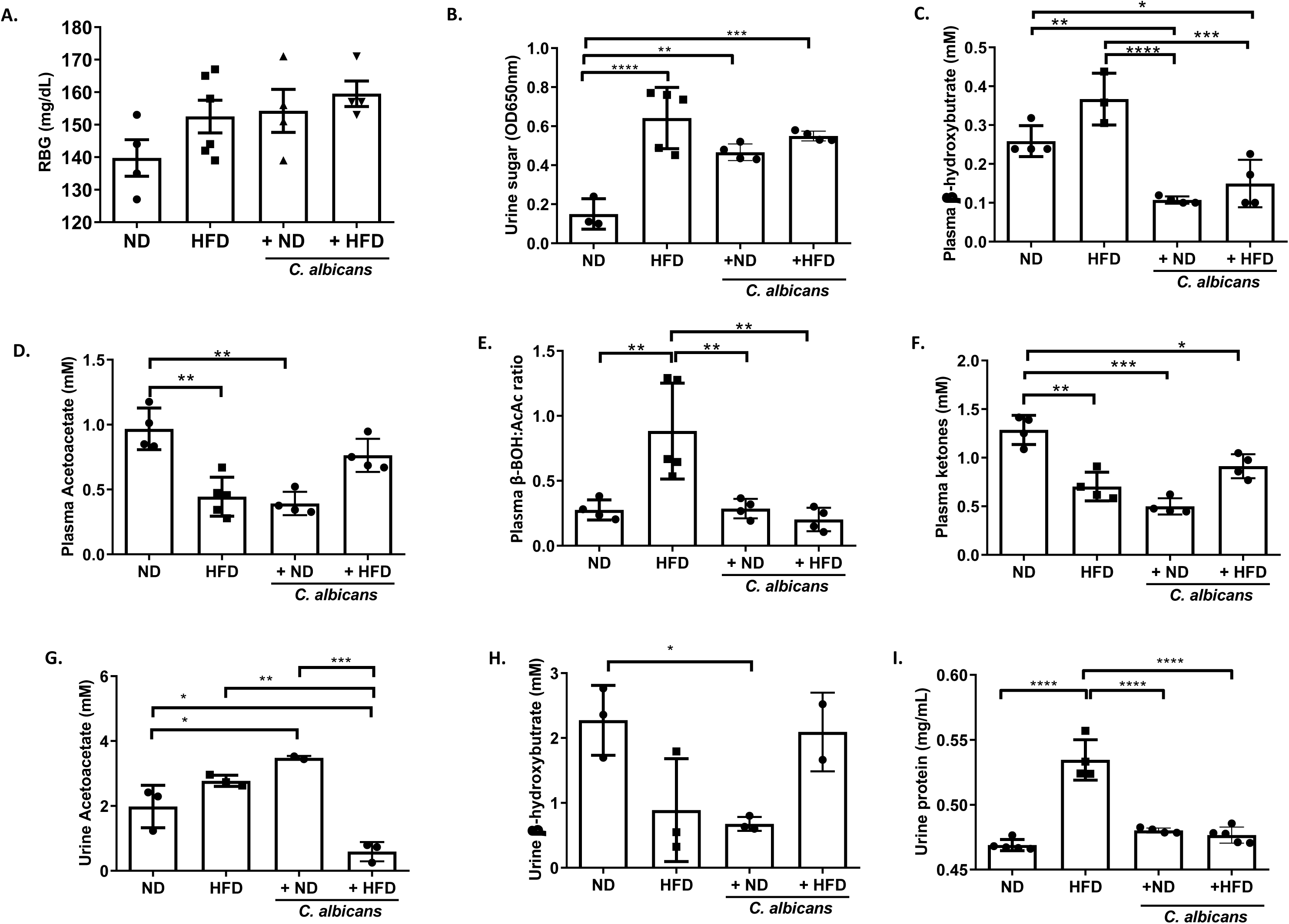
Diet mediated alterations in metabolism. (**A**) Mean and standard error of mean of random blood glucose level (mg/dL), (**B**) urine sugar, (**C**) plasma β-Hydroxybutyrate (mM), (**D**) plasma Acetoacetate (mM), (**E**) β-Hydroxybutyrate to Acetoacetate ratio, (**F**) Plasma Ketones, (**G**) urine Acetoacetate (mM), (**H**) urine β-Hydroxybutyrate (mM), and (**I**) urine proteins (mg/mL) of each mouse from the 4 groups of mice. A statistical significance (*p≤0.05, ** p≤0.01,***p≤0.001, **** p≤0.0001) was calculated using one-way ANOVA and Tukey’s multiple comparison test.

Fatty acids from adipose tissue or a high-fat diet are metabolized in the liver. Under certain circumstances such as carbohydrate restrictive diets, starvation, prolonged intense exercise, alcoholism, low food intake, insulin resistance, or diabetes mellitus, the liver produces ketone bodies by utilizing fatty acids[37]. Acetoacetate, β-hydroxybutyrate (BOH), and acetone are water-soluble molecules collectively called ketone bodies. Here we profiled urinary and plasma ketones to identify HFD induced changes in lipid metabolism as compared to the control diet. The level of plasma BOH was reduced to almost half (0.1–0.15 mM) in the mice fed with any diet when blended with *C. albicans*. In HFD fed mice, the BOH amount was determined to be 0.25 – 0.35 mM **(Fig. 4C)**. In addition, we also found a significant increase in the β-hydroxybutyrate/acetoacetate ratio in HFD that reflects higher BOH levels in the plasma of HFD mice (**Fig. 4C and E**). Surprisingly, the urine excretion of BOH did not reflect the plasma level indicating it’s possible utilization in these mice, however, it was 10 folds higher than that in plasma. The urine BOH levels were high in the mice under normal, and *C. albicans* mixed high-fat diet compared to the other two groups of mice (**Fig. 4H).** The amount of plasma total ketones and acetoacetate in ND fed mice were found to be 1–1.25 mM and that was significantly reduced in HFD and diet with fungal mix (0.5-0.75 mM) (**Fig. 4 D and F**). In general, ketones are not present in urine but appear when fats are burned for energy. About 2–3 mM of acetoacetate was estimated in the urine of HFD fed mice and was drastically reduced to 0.5 mM when the mice were allowed to eat HFD-fungal mix feed. About 2 folds more urine acetoacetate in ND with the fungal mix than the ND alone fed mice was observed (**Fig. 4G**). These results indicated an alteration in glucose and lipid metabolism in HFD fed mice which were restored upon *C. albicans* mix. A high level of plasma BOH is independently associated with cardiovascular events and all causes of death in patients undergoing hemodialysis[38], and the risk seems to be reversed in the presence of *C. albicans* in HFD diet mice.

### Renal functions are altered in HFD but not affected by *C. albicans* feeding

The kidney plays an important role in maintaining metabolic homeostasis through urinary excretion of unwanted and excessive water soluble metabolites. HFD was reported to induce renal injury[39]. Since the kidney is the primary organ for invasive Candidiasis, we evaluated the impact of a high-fat diet and dietary *C. albicans* on renal functions. Although the fungal mixed diet fed mice were healthy and none showed noticeable symptoms of any infections, the kidney size marginally increased in animals feeding fungal mixed diet. An average kidney size in mice on the regular diet was 0.28 gms and it increased maximally to 0.35 gms in mice with fungal mixed HFD (**Supp. Fig. 2B**). To find out any relationship between kidney size and fungal load, the presence or absence of *C. albicans* in kidney cross-sections was examined by Periodic acid–Schiff (PAS) staining. Interestingly, despite 150 days of fungal diet interventions none of the kidney cross-sections showed the presence of *C. albicans*, suggesting that the marginal increase in the size of kidneys in mice fed with fungal mix was not due to the fungal load rather could be due to their overall body weight or high cellular infiltrations in the kidney (**Supp. Fig. 2C**). Also, our results suggested that the impact of dietary *C. albicans* and intravenous challenge of *C. albicans* in mice are very different, and dietary *C. albicans* does not disseminate into the bloodstream to cause any infection. The presence of a urinary protein indicates compromised renal function. We found more proteinuria in the mice fed with HFD (0.53 mg/ml) compared to all other groups of mice (∼0.47 mg/ ml) (**Fig. 4I**). Interestingly urinary protein level in dietary *C. albicans* mixed HFD group was comparable to normal chow without or with fungal mix suggesting that the presence of *C. albicans* in diet improved kidney function in HFD. More importantly, our study also suggested that the gut colonization of *C. albicans* induced by HFD or dietary *C. albicans* is mostly noninvasive.

### Augmented population of circulating immune cells in HFD are restored upon *C. albicans* feed

Cells from both the innate and adaptive immune systems are crucial in the pathogenesis of obesity and its metabolic complications [40]. To understand immune cell homeostasis during experimental obesity induced by HFD and the impact of dietary *C. albicans*, we measured circulating immune cells in the blood (**Fig. 5 A, B, C, and D**). A high level of absolute total WBC, lymphocytes, and monocytes counts were estimated in mice fed with HFD (∼7000 cells/μl, ∼4000 cells/μl, and ∼700 cells/μl respectively) than those in mice fed with normal chow (∼4500 cells/μl, ∼2000 cells/μl, and ∼400 cells/μl respectively). However, the addition of *C. albicans* subdued the effect of HFD and reduced their counts to basal level as in the ND fed mice, additionally, granulocytes counts were also low in *C. albicans* groups. Further based on differential CD11b expression and cell granularity (side scatter, SSC), lymphoid and myeloid compartments in normal chow-fed mice blood were found almost equal percentage (each ∼45-55% of total PBMC). However, a significantly high level of circulating lymphoid cells (∼65-70 %) than the myeloid cells (∼35 %) was found in the HFD alone and *C. albicans* mixed ND groups (**Fig.5 E, F, and G**). Interestingly, *C. albicans* addition in HFD fed restored the lymphoid and myeloid distribution to the normal chow level (∼45 % and 55 % respectively). These data suggest that dietary *C. albicans* impact on the immune system might be dependent on dietary calorie and host physiology. Further high circulating lymphoid compartment along with a high absolute count of immune cells such as lymphocytes and monocytes in the HFD fed mice per se mimics inflammatory status quo as found in infections. Importantly, these changes were found to be restored upon *C. albicans* presence in HFD.

**Figure 5.**
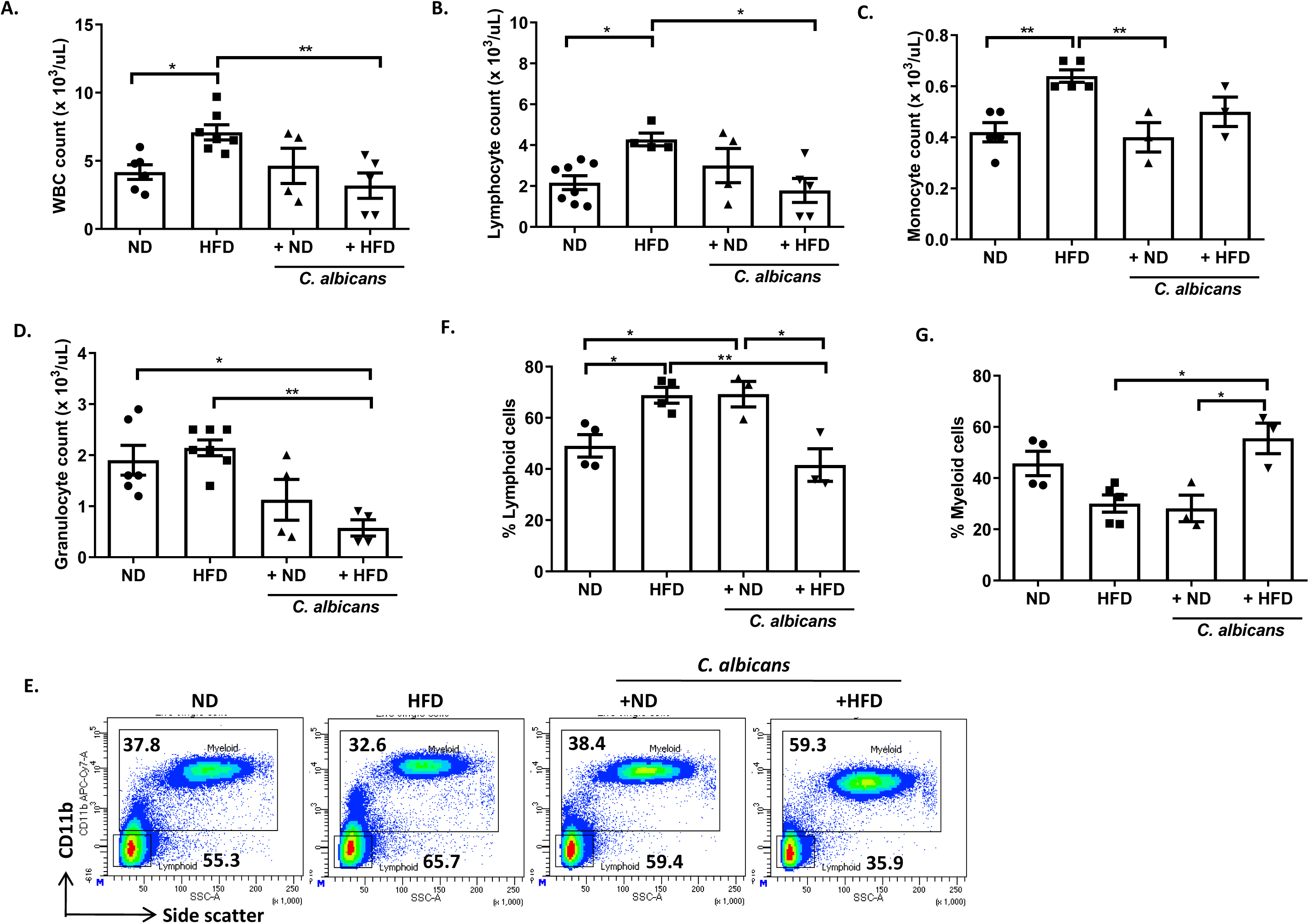
Changes in peripheral blood circulating immune cells. Mean and standard error of mean of absolute (**A**) WBC, (**B**) Lymphocyte, (**C**) Monocyte, and (**D**) Granulocyte counts from blood of all the 4 groups mice on 150th day of dietary intervention. (**E**) A representative bivariate density plot of CD11b (y- axis) vs Side scatter (x-axis) for the analysis of percent total Myeloid and Lymphoid population of blood cells using BD FACSDiva Software v8.0.2 . Mean and standard error of mean of percent total (**F**) Lymphoid cells, and (**G**) Myeloid cells from peripheral blood of all the 4 groups mice on sacrifice (150th day), a statistical significance (*p≤0.05, ** p≤0.01,***p≤0.001, **** p≤0.0001) was calculated using one-way ANOVA and Tukey’s multiple comparison test.

### *C. albicans* supplemented diet restored HFD induced high circulating myeloid macrophages and autoimmune prone CD5^+^B cells to normal level

For better insights into the differential cellular composition of immune cells in circulation, cells were stained for multicolor immune phenotyping and analyzed by gating based on their specific markers (**Fig. 6A I and B I**). To characterize myeloid cells in the blood, we considered cellular population with CD11b^+^/Gr-1^+^hi/SSC(hi) as neutrophil, CD11b^+^/Gr-1^+^/SSC(lo) as monocyte, CD11b^+^/Gr-1^+^/SSC(hi) as eosinophil, and CD11b^+^/Gr-1-/SSC(lo) as myeloid dendritic cells & macrophages **(Fig. 6A I)**. Percentage of monocytes, neutrophil, and eosinophil are more or less remain unaltered in various mice groups except for a minor reduction of monocyte in mice with HFD-fungal mixed feed (∼ 1.5 – 3 %) (**Fig. 6A II**). However, the percentage of circulating macrophages and myeloid dendritic cells population increased to 12.5 % in mice fed with a high-fat diet in comparison to 8 % in mice with a regular diet. Again, the supply of dietary *C. albicans* substantially reduced their populations to ∼ 2-3 % irrespective of diets intake (**Fig. 6A III**). Lymphoid populations were characterized as T cells (CD11b^-^/CD19^-^/CD5^+^), B Cells (CD11b^-^/CD5^+^/^-^/CD19^+^) and non-T or B lymphoid cells, such as NK, lymphoid DCs (CD11b^-^/CD5^-^/CD19^-^). Surprisingly, the T cells population (∼40 % Vs ∼20 %) was increased in the peripheral circulation of mice fed with *C. albicans* mix irrespective of diet content than the mice without fungal addition **(Fig. 6B II**). CD5 is actually a T cell marker that is expressed in a subset of B cells as well. CD5^+^ B cell subtypes in circulating lymph were estimated to be ∼14 % in the ND fed mice that were drastically reduced (2.5 –6 %) in mice when fed with a *C. albicans* mixed diet. A noticeable increase in CD5^+^ B cell types (17.5 %) in HFD alone group mice was observed **(Fig. 6B III)**. An increased population of circulating macrophages, dendritic, and CD5 positive B cells in HFD fed mice suggests possible inflammation induced by obesity or overweight, and the presence of *C. albicans* suppresses the impacts of HFD. Although an increase in circulating T cells is usually found in infection, the increased population of T cells in fungal mixed over regular or HFD fed mice that had no features of comorbidity or fungal load in vital organs indicates a strong impact of gut microbes on altering circulating T cells.

**Figure 6.**
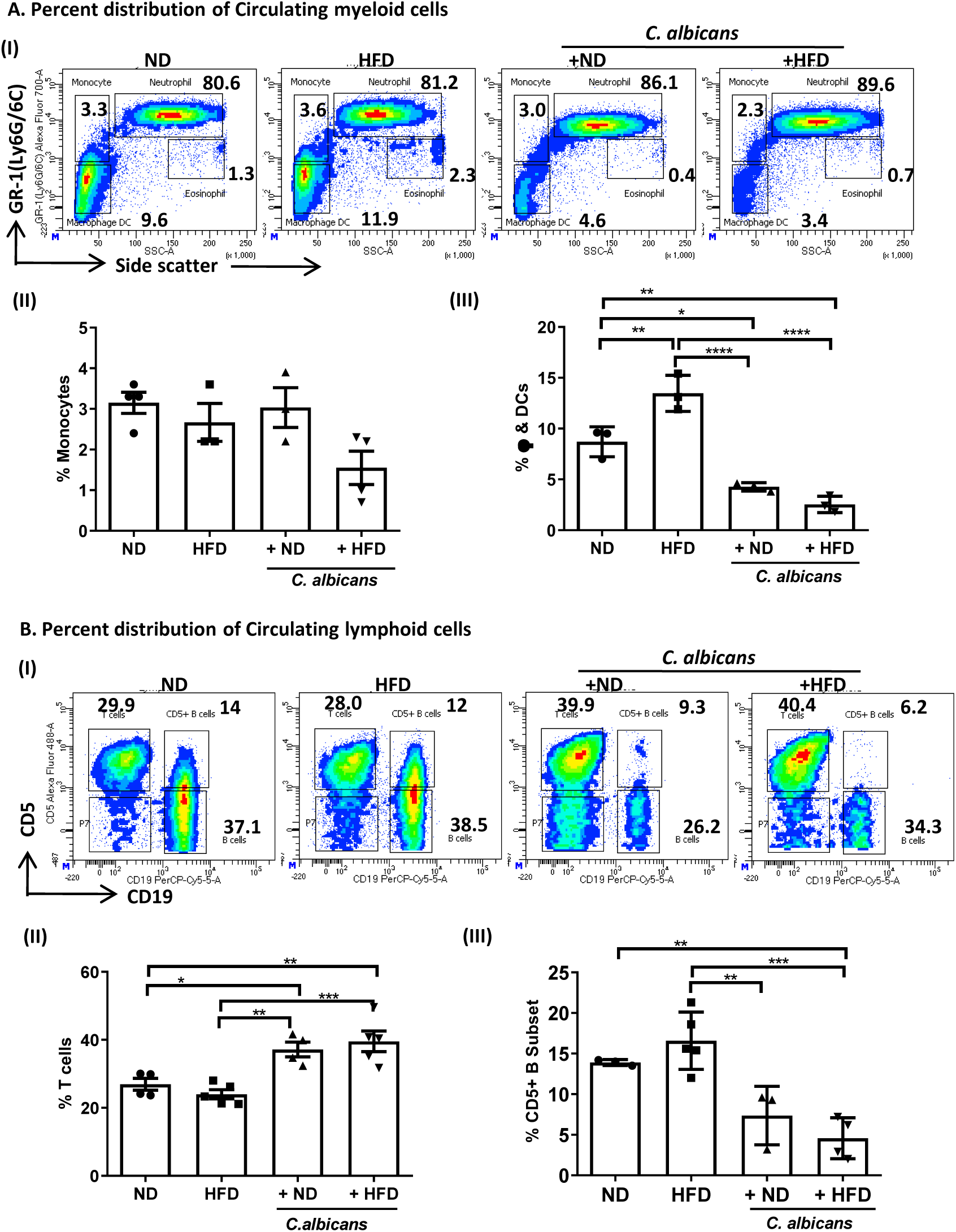
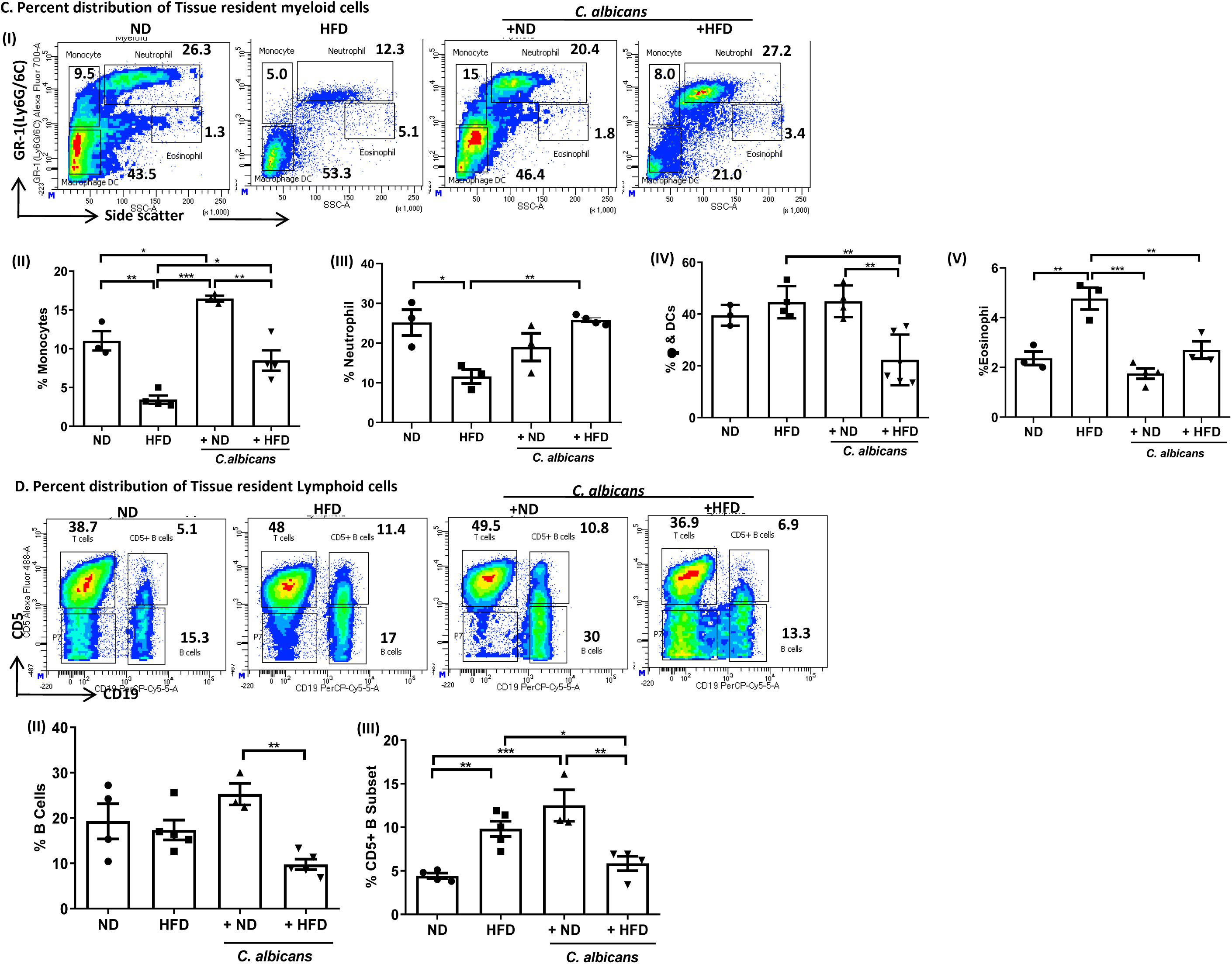
Distribution of circulating and tissue resident myeloid and lymphoid compartmental immune cells, (**A (I)**) A representative bivariate density plot of Gr-1(Ly-6G/Ly-6C) (y-axis) vs Side scatter (x-axis) for the analysis of compartmental distribution of CD11b^+^ Myeloid cells using BD FACSDiva Software v8.0.2. Mean and standard error of mean of percent total (**A (II)**) Monocytes, and (**A (III)**) Myeloid macrophages & Dendritic cells of peripheral CD11b^+^ Myeloid compartment from blood of all the 4 groups mice on the day of sacrifice. (B **(I)**) A representative bivariate density plot CD5 (y-axis) vs CD19 (x-axis) of lymphoid population for the analysis of their subpopulation using BD FACSDiva Software v8.0.2. Mean and standard error of mean of percent total (**B (II)**) Thymic lymphocytes, and (B **(III)**) CD5^+^ B lymphocytes of peripheral lymphoid compartment from blood of all the 4 groups mice on 150th day. (**C (I)**) A representative bivariate density plot of Gr-1(Ly-6G/Ly-6C) (y-axis) vs Side scatter (x-axis) for the analysis of compartmental distribution of CD11b^+^ Myeloid cells in spleen using BD FACSDiva Software v8.0.2. Mean and standard error of mean of percent total (**C (II)**) Monocytes, (**C (III)**) Neutrophil (**C (IV)**) Myeloid macrophages & Dendritic cells and (**C (V)**) Eosinophils of tissue resident CD11b^+^ Myeloid cells in spleen of all the 4 groups mice on the day of sacrifice. (**D (I)**) A representative bivariate density plot CD5 (y-axis) vs CD19 (x-axis) of lymphoid population for subpopulation analysis using BD FACSDiva Software v8.0.2. Mean and standard error of mean of percent total (**D (II)**) B lymphocytes, and (**D (III)**) CD5^+^ B lymphocytes of tissue resident lymphoid compartment in spleen of all the 4 groups mice on the day of sacrifice. A statistical significance (*p≤0.05, ** p≤0.01,***p≤0.001, **** p≤0.0001) was calculated using one-way ANOVA and Tukey’s multiple comparison test.

### HFD modulate splenic tissue resident immune cells distribution that returned to normal upon *C. albicans* supplementation

Lymphoid organ resident immune cells initiate immune responses and restore homeostasis during any infection, stress, and other deviations from the normal[41]. Since aberrant circulating immune cell profile was observed in the HFD fed mice, we further sought to understand the immune cell distribution in secondary lymphoid organs, such as in the spleen. On contrary to circulating immune cell profile, based on CD11b expression and internal granularity, the tissue resident proportion of myeloid cells (∼ 30%) were less than the lymphoid populations (∼ 50%) in the spleen. The myeloid population was further reduced in HFD (∼ 20%), but a noticeable increase of (∼ 38%) in mice fed with HFD mixed with *C. albicans* culture, however, no change was found among the other groups of the cohort (**Supp. Fig. 3 I and II**). Lymphoid cells population increased in HFD consumed mice (∼75 %) than that in ND-fed mice (∼50-60 %), again it was controlled by *C. albicans* dietary intervention (∼40 %) (**Supp. Fig. 3 III**). Based on of GR-1 expression and granularity, myeloid populations were further segregated to Monocyte, Neutrophil, Eosinophil, and myeloid macrophage DCs lineages. Splenic monocytes and neutrophil populations decreased in the mice fed with an obesogenic diet (2.5 % and 11 %, respectively) than the mice with a normal diet (11 % and 25 %, respectively). By the supplementation of *C. albicans* cells to HFD, it almost restored to the basal level of the tissue resident monocytes and neutrophil populations (9 % and 25 %, respectively). We observed a marginal increase of monocytes (17 %) but no change in neutrophils in the spleen of mice consuming fungal mixed regular diet (**Fig. 6C I, II, and III**). Splenic macrophages and DCs did not alter among the mice groups (∼40 %) except in the mice consuming fungal mixed HFD (∼18 %) (**Fig. 6C I, IV**). Surprisingly, the eosinophil populations almost doubled in mice fed with obesogenic food (5 %) than that in regular diet (2.5 %), however, the presence of *C. albicans* in the diet helped in maintaining the eosinophil population (**Fig. 6C I and V**). Lymphoid populations that include T cells, B cells, and non-T or B lymphoid cells, such as NK, lymphoid DCs were analyzed among tissue residents. Fatty diet appeared not to affect tissue resident total B cells population (∼20 %), however, the presence of fungal cells only in the HFD diet marginally reduced the total B cells percentage (9 %) (**Fig. 6D I and II**). CD5^+^ B cell subtypes increased 2-3 folds in mice (10-12.5 %) with HFD alone or *C. albicans* supplemented normal chow than the control mice (∼5%). But the addition of *C. albicans* to HFD restored the basal level CD5^+^ B cell population (**Fig. 6D I and III**). This observation was significant and consistently similar on both tissue resident and circulating CD5^+^ B cells in the groups.

### Dietary factors induced differential CD4^+^ T cell immune response

Our results have indicated a possible compromised state of cellular immune response in HFD fed cohort that could be improvised on supplementation of dietary *C. albicans.* CD4^+^ T cells play an important role in the progression of obesity and obesity-associated disorders [42]. Circulating CD4^+^ T cell frequency is positively associated with increased body mass index (BMI) or obesity conditions in humans[43]. To identify the adaptive CD4^+^ T cells response to dietary *C. albicans* and HFD, we have isolated CD4^+^ T cells from the spleen of all four groups of mice and stimulated them *in vitro* to characterize their cytokine and immune inhibitory response status. We found that TNF-α^+^ CD4^+^ and INFγ^+^ TNFα^+^ CD4^+^ T cells population doubled when mice were subjected obesogenic diet in comparison to the normal diet fed mice (**Fig. 7 A, C, and D)**. However, ∼70 % reduction of INFγ^+^ CD4^+^ and CTLA4^+^ CD4^+^ T cells population were estimated in mice fed with HFD in comparison to the regular diet fed mice (**Fig. 7 B, E, and G)**. Concomitantly, diet preference did not affect the population of IL17A^+^ CD4^+^ and IL17A^+^ CTLA4^+^ CD4^+^ T cells (**Fig. 7 E, F, and H)**. The addition of *C. albicans* cells to the diet had a broad range of effects. Irrespective of the food intake, presence of *C. albicans* lowered the percentage INFγ^+^ CD4^+^, INFγ^+^ TNFα^+^ CD4^+^, CTLA4^+^ CD4^+^, and IL17A^+^ CTLA4^+^ CD4^+^ T cells population (**Fig. 7 B, D, G and H)**. Only, a high percentage of TNFα^+^ CD4^+^ and IL17A^+^ CD4^+^ T cells population were found in mice fed with the *C. albicans* supplemented diet (**Fig. 7 C and F).** The high percent CTLA4^+^ CD4^+^ T cells along with low percent INFγ^+^ CD4^+^ and INFγ^+^ TNFα^+^ CD4^+^ T cells with no IL17A in normal chow indicated the mice to have a well-regulated and controlled immune status. However, low percent CTLA4^+^ CD4^+^ T cells along with high percent INFγ^+^ CD4^+^ and INFγ^+^ TNFα^+^ CD4^+^ T cells with no IL17A in the HFD group of mice suggested uncontrolled inflammatory conditions that could predispose the animals to autoimmunity and inflammatory diseases.

**Figure 7.**
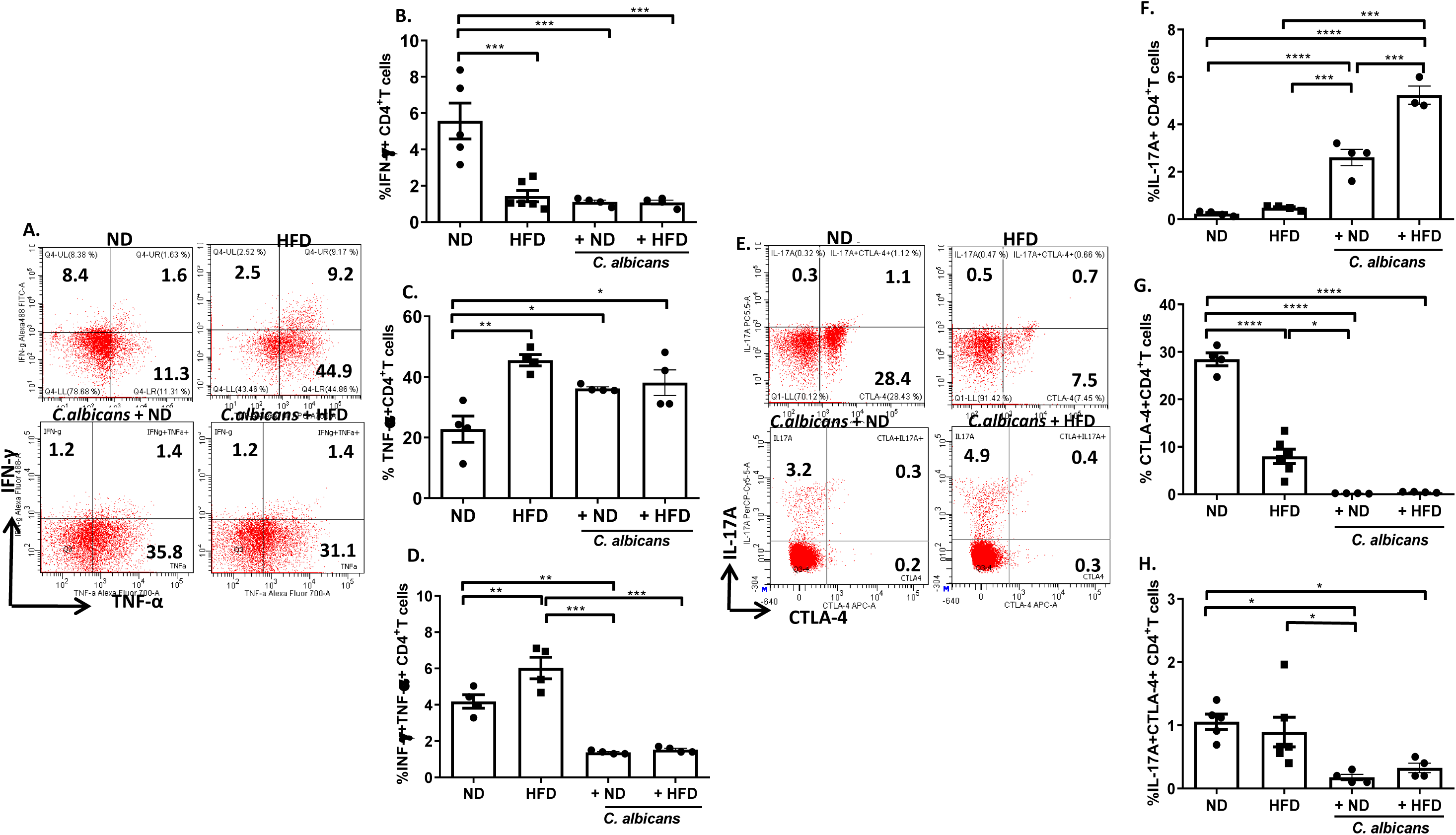
Alterations in Tissue resident CD4^+^ T cells responses occurred upon dietary manipulation. A representative bivariate density plot of (**A**) IFNγ^+^ (y-axis) vs TNFα^+^ (x-axis) and (**E**) IL17A^+^ (y-axis) vs CTLA-4^+^ (x-axis) of CD4^+^ T lymphocytes for functional analysis using CytExpert Software for above two images ND and HFD acquired in Cytoflex and BD FACSDiva Software v8.0.2 was used for below two images *C. albicans* + ND and *C. albicans* + HFD acquired in BD LSRFortessa. Mean and standard error of mean of percent CD4^+^ T cells positive for (**B**) IFNγ^+^, (**C**) TNFα^+^, (**D**) Double IFNγ^+^ TNFα^+^ population for all four groups of mice, (**F**) IL17A^+^, (**G**) CTLA4^+^, and (**H**) Double IL17A^+^ CTLA4^+^ populations analyzed for all four groups of mice. A statistical significance (*p≤0.05, ** p≤0.01,***p≤0.001, **** p≤0.0001) was calculated using one-way ANOVA and Tukey’s multiple comparison test.

Interestingly, anti-fungal IL-17 response seems to be intact in both the diets supplemented with *C. albicans* irrespective of the obese status, whether the activation of a similar immune response to fungal infection after the onset of obesity necessitates further investigation.

### *C. albicans* infection challenge revealed compromised anti-fungal CD4^+^ T cell responses in HFD fed mice

Since the HFD fed mice without *C. albicans* supplementation exhibited uncontrolled inflammatory profiles compared to a well-regulated and controlled immune status in a normal diet fed mice, further, we determined their immune response to the intraperitoneal infective challenge of *C. albicans.* After induction of DIO with 21 weeks of feeding with HFD or ND, the mice were subjected to IP challenge of the fungal pathogen and after 48 hrs, animals were sacrificed, splenic CD4^+^ T cells were isolated, collected, and monitored immune response by in vitro stimulation. We found a higher population of TNFα^+^ CD4^+^ (∼ 45 % *Vs* 21 %) and INFγ^+^ TNFα^+^ CD4^+^ T (∼7 % *Vs* 4.5 %) cells in mice subjected to the obesogenic diet in comparison to the normal diet fed mice (**Fig. 8 A, C. and D)**. However, ∼ 80 % reduction of INFγ^+^ CD4^+^ cells population was estimated in mice fed with HFD in comparison to the regular diet fed mice (**Fig. 8 A and B**). At the same time, the IP challenge did not affect these populations, unlike fungal mixed diets. Contrastingly, CTLA4^+^ CD4^+^ T cells population increased by IP challenge in the HFD fed mice, and an opposite trend was found in mice fed with regular diet (**Fig. 8 E and G**). Interestingly, the IL17A^+^ CD4 population was very low in HFD fed mice with and without IP challenge, and a good IL17A immune response to the IP challenge was observed in mice subjected to normal food (**Fig. 8 E and F)**. No significant difference in double positive IL17A^+^ CTLA4^+^ CD4^+^ T cells was observed in these sets of mice (**Fig. 8 E and H**). Irrespective of the routes of *C. albicans* intervention i.e. as a food or IP challenge, the mice elicit IL17A immune response when they were subjected to a non-obesogenic diet but not HFD. Additionally, these results suggested that regardless of dysregulated inflammatory status in HFD mice that could succumb and not be able to mount an immune response against infectious fungal challenge pointing to compromised immunity in obesity.

**Figure 8.**
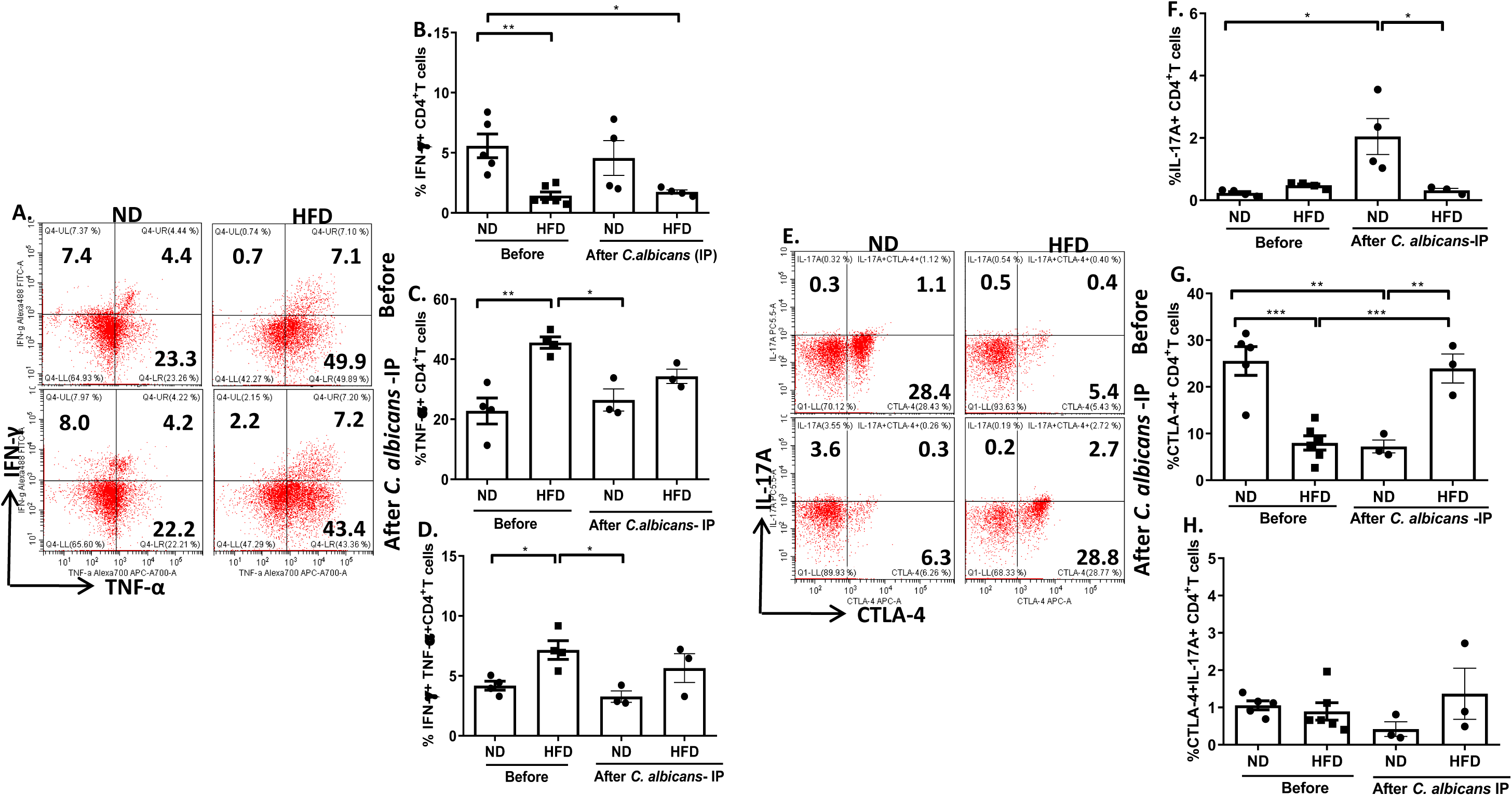
A Comparative tissue resident CD4^+^ T cells responses in ND and HFD fed mice upon *C. albicans* intra peritoneal challenge. A representative bivariate density plot of (**A**) IFNγ^+^ (y-axis) vs TNFα^+^ (x-axis) and (**E**) IL17A^+^ (y-axis) vs CTLA4^+^ (x-axis) of CD4^+^ T lymphocytes for functional analysis using CytExpert Software comparing ND and HFD before (above) and after (below) intra peritoneal *C. albicans* challenge. Mean and standard error of mean of percent CD4^+^ T cells positive for (**B**) Interferon gamma^+^ (IFNγ^+^), (**C**) Tumor necrosis factor- alpha^+^ (TNFα^+^), (**D**) IFNγ^+^ TNFα^+^ population, (**F**) Interleukin-17A^+^ (IL17A^+^), (**G**) Cytotoxic T-lymphocyte-associated protein 4^+^ (CTLA4^+^), and (**H**) Double IL17A^+^ CTLA4^+^ populations analyzed for ND and HFD before (above) and after (below) intra peritoneal *C. albicans* challenged mice, A statistical significance (*p≤0.05, ** p≤0.01,***p≤0.001, **** p≤0.0001) was calculated using one-way ANOVA and Tukey’s multiple comparison test.

### HFD and *C. albicans* differentially regulate gut microbiota

As the microbiome plays an important role in the health status of an individual and the observation that dietary *C. albicans* colonizes in the gut (**Fig. 1D**), we wanted to monitor the effect of a high-fat diet and exogenously added *C. albicans* on the diversity and richness of gut microbiota. As the gut is mostly filled with bacteria and fungi, we used the metagenomics approach to identify various microbes by sequencing of 16S rDNA and Internal Transcribed Spacer (ITS) region of 18S rDNA from fecal meta-DNA samples (**Suppl. Fig. 4, Fig. 9**). Our 16S rDNA analyses revealed an abundance of 1263 to 1695 bacterial species in all four samples (**Suppl. Table 2**). HFD fed mice showed poor microbial richness (1263 Operational taxonomic units (OTUs)), however mixing of dietary *C. albicans* with HFD restored the microbial composition similar to in mice fed with normal diets (1500-1695 OTUs). Thus, while HFD induces dysbiosis, dietary *C. albicans* has a positive influence on gut microbial diversity. Alpha rarefaction plot that helps in comparing diversity across samples also indicated poor richness of microbial species in HFD fed mice (**Suppl. Fig. 5A**). Among the four samples, only 112 bacterial species were found to be common, and 1134, 1189, 825, and 1366 species were specifically detected in the guts of mice fed with a normal diet, a normal diet with *C. albicans* mix, a high-fat diet, and a high-fat diet with *C. albicans* mix, respectively (**Fig. 9A**). Less than 1 % of sequences corresponding to unclassified or unknown bacteria were detected in each sample. The proportion of bacteria from Firmicutes and Bacteroidetes contribute maximally to healthy gut microbiota. Obese individuals were reported to possess relatively decreased Bacteroidetes and increased Firmicutes in their guts when compared to lean subjects, and in this study also, a similar trend was observed involving Balb/c mice as a model organism. Interestingly, although the alteration in richness proportion of these two bacterial phyla was quite apparent between normal and high-fat diet samples, we found that the percentage of Firmicutes (∼33-35 % in normal diet samples Vs 53-59 % in HFD samples) and Bacteroidetes (∼51-53 % in normal diet samples Vs 16-27% in HFD samples) among the samples did not alter significantly by the dietary intervention of *C. albicans* **(Fig. 9B).** In addition to these two phyla, species belonging to Proteobacteria were the third abundant species, and they were mostly enriched in all the samples (∼8-11 %) except in normal diet fecal samples where they were poorly detected (∼1 %). Thus, Proteobacteria could help in gaining healthy body weight. Further, the OTUs were analyzed to distribute them in the top 20 classes, orders, and families (**Suppl. Fig. 5 B, C, and D).** At the genus and species level, the addition of fungal mix to the diet modulated the bacterial composition drastically **(Fig. 9 C and D).** For example, *Lactobacillus* species were highly abundant in HFD but reduced in the presence of *C. albicans* (64 % to 15 %), and the effect was reversed in normal diet samples (increased from 20 to 60 % with dietary *Candida*). Our metagenomics analyses also revealed the presence of diet specific bacteria. The normal diet fed mice only possessed *Pedicococcus* sp., while *Streptococcus* and *Staphylococcus* species were only detected in the fungal mixed high-fat diet **(Suppl. Table-3**).

**Figure. 9.**
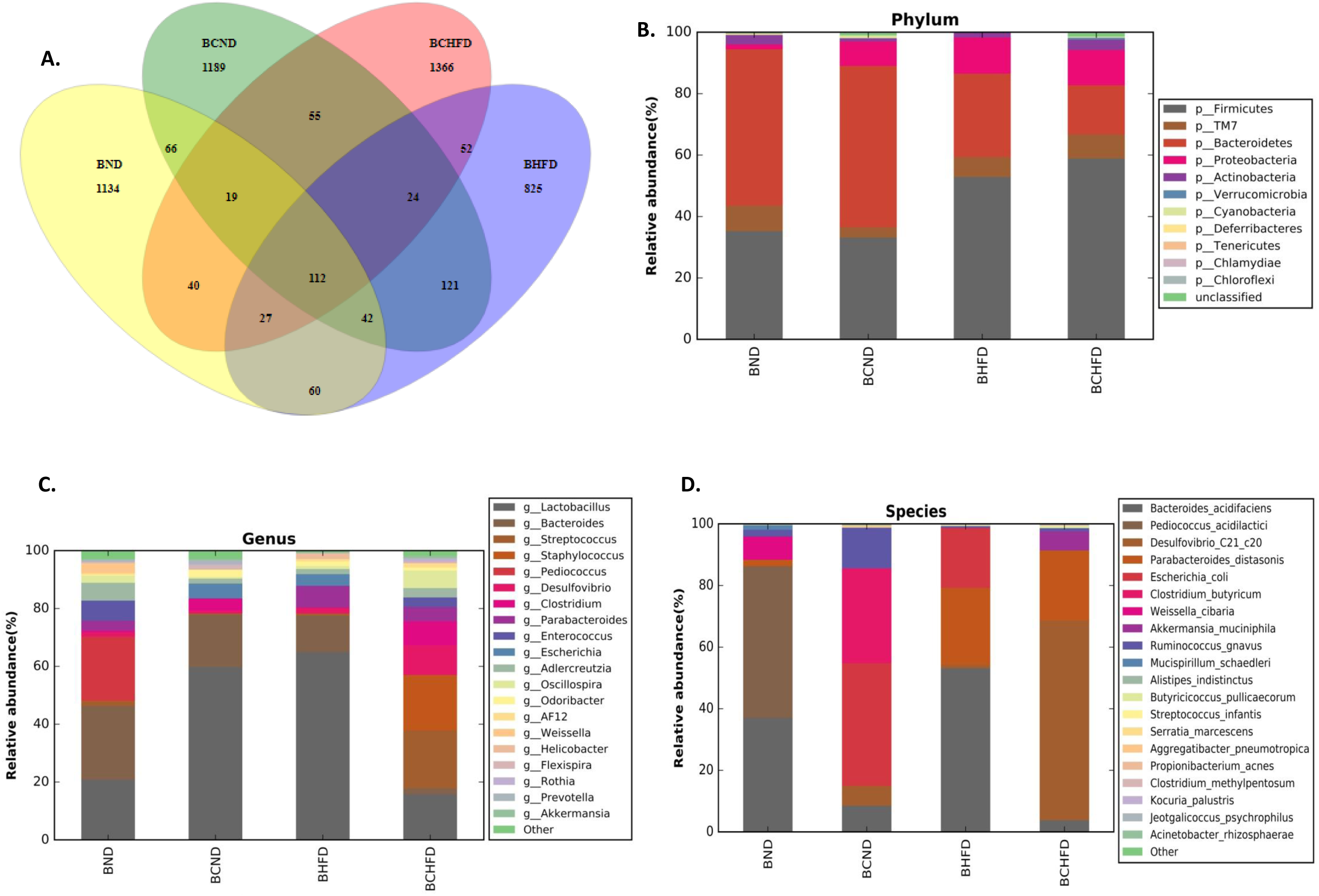
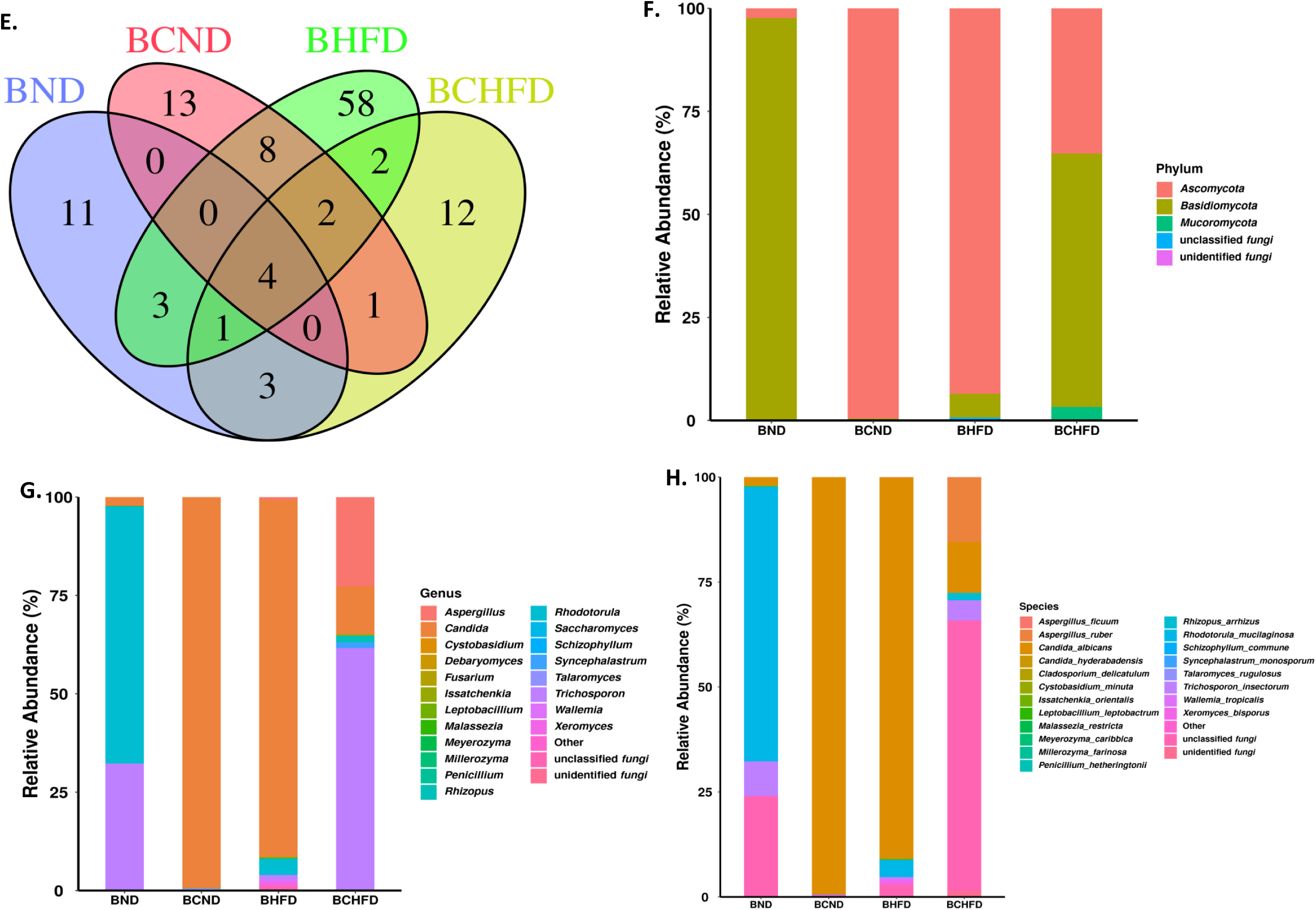
16s rDNA and 18s rDNA metagenomics analyses. A Venn diagram depicting unique and shared OTUs among the four cohorts **(A)**. Taxonomic classification was carried out and assigned them into top 10 phylums (**B**), top 20 genus (**C**) and top 20 species (**D**). A Venn diagram depicting unique and shared ASVs among the four cohorts **(E)**. Taxonomic classification was carried out and assigned them into top 10 phylums (**F**), top 20 genus (**G**) and top 20 species (**H**). BND, metagenomic DNA isolated from normal diet fed BALB/c mice fecal sample; BCND, metagenomic DNA isolated from normal diet with *C. albicans* mix fed BALB/c mice fecal sample; BHFD, metagenomic DNA isolated from high-fat diet fed BALB/c mice fecal sample; and BCHFD, metagenomic DNA isolated from normal diet with *C. albicans* mix fed BALB/c mice groups fecal sample.

The Internal Transcribed Spacer (ITS)-based sequencing effort yielded 205305-267408 sequence reads, of which 47840-96481 corresponded to 118 amplicon sequence variants (ASVs) belonging to 3 different phyla, 11 classes, 18 orders, 28 families, and 39 different genera of fungi (**Suppl. Table-2, Fig. 10**). Fungal sequences were detected in every sample analyzed. Less than 1% of sequences corresponding to unclassified or unknown fungi were detected in each sample. Interestingly, only 4 ASVs were common to all the four samples sequenced, and according to the alpha diversity index plot DNA sample of *C. albicans* mixed HFD showed the maximum variation (**Fig. 9 E and Suppl. Fig 6A)**. Fungi belonged to the phyla Ascomycota and Basidiomycota were abundant in all the samples. Fungal species belonging to the phylum Basidiomycota (97.6%) were enriched in normal died fed mice gut, whereas HFD fed or *C. albicans* mixed died fed mice showed an abundance of fungi belonged to Ascomycota (∼93.5 – 99.5 %). Interestingly, the presence of *C. albicans* in the HFD alters the ratio of Ascomycota and Basidiomycota from 16:1 (93.54 % Vs 5.74%) to 0.6:1 (35.25 % Vs 61.51 %) in the gut, also fungi belonged to Mucoromycota got enriched (**Fig. 9F**). While species belonging to classes Saccharomycetes, Sordariomycetes, and Tremellomycetes were commonly found in all the samples, dietary *C. albicans* suppressed enrichment of Cystobasidiomycetes, Dothideomycetes, Microbotryomycetes classes of fungi. The abundance of Eurotiomycetes and Mucoromycota were only found in HFD fed mice samples, but the addition of *C. albicans* to the food reduced their composition significantly (**Suppl. Fig 6 B, C, and D**). At the genus level, *Candida* and *Trichosporon* fungi were commonly found in all the samples, whereas *Aspergillus* and *Fusarium* were observed in all the samples except control diet fed mice. Additionally, we also observed diet specific abundance of the fungal genus (**Fig. 9 G and H)**. While in the normal diet fed mice only *Monographella, Neoascochyta, Papiliotrema*, and *Wickerhamomyces* fungus were enriched, species belonging to *Acremonium, Cladosporium, Clavispora, Didymella, Leptobacillium, Issatchenkia, Microascus, Millerozyma, Myceliophthora, Ovatospora, Penicillium, Saccharomyces, Starmerella, Talaromyces,* and *Xerochrysium* genus were seen in HFD subjected mice. Similarly, *Naganishia, Schizophyllum*, and *Sterigmatomyces* genus and *Mucor, Rhizopus*, and S*yncephalastrum* genre were abundant in *C. albicans* mixed normal and high calorie content diet, respectively (**Suppl.Table-3**). Consistent with the genus level, *C. albicans* species were the standout species in all the samples. Other species such as *C. dubliniensis*, *C. hyderabadensis*, and *C. sake* were also observed in a few samples. To our surprise, we did not detect other human prevalent *Candida* species such as *C. tropicalis*, *C. parapsilosis, C. glabrata, C. krusei,* and *C. lusitaniae*[14, 44]. As shown in Fig. 1D as well, low but significant abundance (∼2%) of *C. albicans* was found in the metagenome of normal died fed mice, but the addition of dietary *C. albicans* to the normal diet or HFD alone facilitated its better colonization (∼99 -90 %). Surprisingly, only a marginal increase in the colonization of *C. albicans* (12 %) in the gut of *C. albicans* mixed HFD fed mice was noticed. This result suggests that while HFD by itself facilitates *C. albicans* growth in the gut, the presence of dietary *C. albicans* somehow negates the effect of HFD on its own colonization. In terms of fungal species composition, HFD metagenome is highly diversified and 28 different species of fungus were seen. However, the addition of *C. albicans* to HFD reduced the species diversity drastically to only 10 different species. Fungal species diversity remained low in the normal diet fed mice irrespective of dietary *C. albicans* addition. *Rhodotorula mucilaginosa* and *Trichosporon insectorum* were the highly enriched species in control diet fed mice metagenome. *Aspergillus ruber* and *Rhizopus arrhizus* are the two notable fungal species that got specifically enriched by the addition of dietary *C. albicans* to HFD. From our metagenomics analyses, we conclude that dietary *C. albicans* colonizes in the host and regulates its own colonization in the host consuming a high-fat diet, in addition to regulating other microbial species abundance. Diet types play an important role in determining the gut microbial composition of both bacterial and fungal species, obesity, and immunity, even in the BALB/c mice model. Alternations in the gut microbiota could influence the metabolites released by bacterial and fungal species which are known to impact host physiology and immunity [45–47], thus, *C. albicans* being a commensal plays an important role in maintaining eubiosis and suppressing complications aroused due to HFD (**Fig. 10**). Microbial metabolites like butyrate, lipopolysaccharides (LPS), Short-chain fatty acids (SCFAs) were found to regulate obesity, diabetes, bone development etc [48].

**Figure 10.**
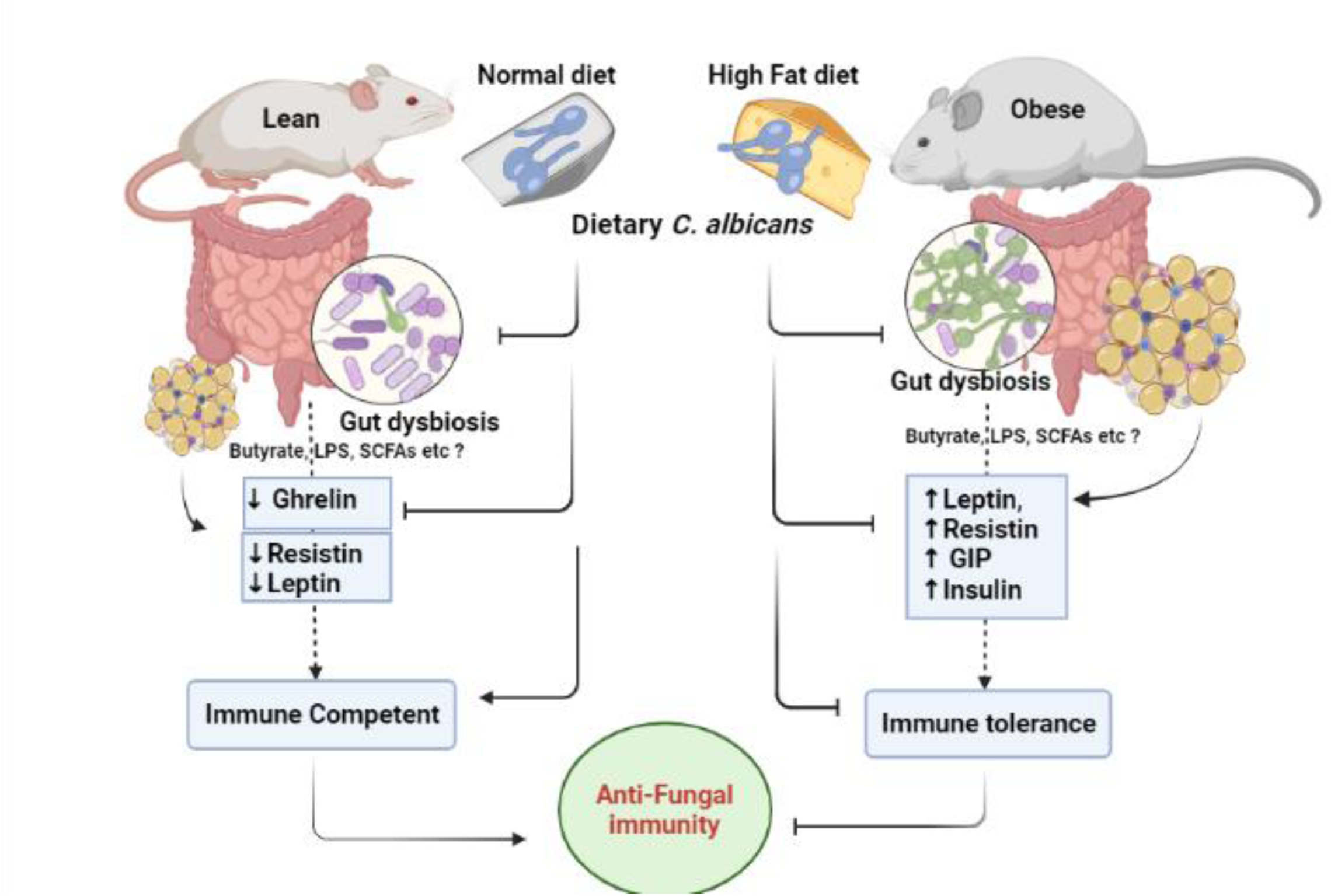
A possible mechanism of dietary *C. albicans* affecting obesity and associated outcomes in mice. A holistic representation of the effect of dietary *C. albicans* as a probiotic in modulating gut microbiome of mice which in turn release different metabolites in different diet conditions. Metabolites may bind to various tissue specific receptors to alter metabolic parameters, gain in body weight, immune cells profiles, and immunity in mice (curtsey to BioRender).

## Discussion

For decades it is known that *C. albicans* has no known terrestrial life cycle, it only survives and co-exists both in humans and animals as a commensal organism. Several studies have been carried out considering it only as a pathogenic yeast but any beneficial symbiotic relationship that may exist between the host and the fungi was not explored until the present study. A cooperative interaction between the microbiome and its hosts is critical in modulating host defense, metabolism, and reproduction [12, 45, 49, 50]. Distinct microbial compositions were found in lean and obese subjects, although limited studies are available to decipher the precise role of individual bacterial and fungal species in the host’s functions. Many reports have suggested that *C. albicans* gut load is associated with obesity regardless of other microbes involvement [17, 51, 52]. Interestingly a diet that controls *Candida* growth in the gut, the “anti-Candida diet” has also been on prescription for preventing obesity without any clear piece of evidence that supports the practice [53–56]. Importantly since one of the sources of gut microbe load has been the diets, we have attempted here to understand the role of dietary *C. albicans* in obesity, associated metabolic and immunological outcomes, and its influence on gut microbial composition by using BALB/c as the model organisms. Since the diet-induced obesity mice models mimic human metabolic abnormalities developed during obesity and share the gut microbial ecology, they become the obvious choice of investigations[57]. Several groups have shown that about 12-22 weeks of high calorie content food consumption results in ∼40-80% body weight gain and promoted obesity in various strains of mice [21, 58]. Herein, we found that HFD consumption in BALB/c mice gained ∼50 % body weight of its original in about 22 weeks, which is 2 folds higher than the mice fed with normal chow for the same duration. Thus, we argued that the BALB/c strain is not a DIO resistant strain as previously reported, only it takes a longer feed duration (22 weeks) to gain body weight compared to other mice strains (12-16 weeks). Nevertheless, despite differences in kinetics, most of the mice strains develop obesity by continuous feeding of a high-calorie content diet. Interestingly and contrary to the practice of anti-Candida diet use, *C. albicans* presence in the diet could reduce the excessive body weight gain, the effect was not either due to infection as the mice were active and healthy and or due to low food intake. Th1 immunity is activated during colonization of *C. albicans* in mice, and the development of acquired resistance is associated with strong Th1 responses [59]. Although the BALB/c mice are susceptible to disseminated systemic candidiasis as they are Th2 biased strain [60], dietary *C. albicans* did not cause any noticeable infection irrespective of the types of diets consumed by the mice. It also indicates that overgrowth of *C. albicans* is not sufficient to cause fungal infection and it may require some host factors. In fact reports suggested that dysbiosis in the gut is not sufficient to promote the invasive growth of *C. albicans*, additionally, it requires immunosuppression of the host and damage on the mucosal layer [61, 62]. A summary of our findings related to *C. albicans* modulating various parameters of obesity related pathological disorders in mice has been elucidated in **Suppl. Fig. 7**.

### Metabolic alterations by HFD and its mitigation by dietary *C. albicans*

Our results revealed alterations in multiple biochemical parameters and hormonal imbalance. HFD fed mice were with excessive abdominal fat deposits, hyperinsulinemia, glycosuria, ketonuria, proteinuria, and imbalances of metabolic hormones and appetite-regulating neuropeptides. These alterations however seem to be regulated and restored by the gut load of *C. albicans* in BALB/c mice via feeding. Release of increased leptin but reduced ghrelin hormones by HDF fed mice clearly suggested imbalance in food intake versus consumption in these mice, and reversal of this effect was observed in *C. albicans* blended diet. Similar to our study, the leptin/ghrelin ratio was found to be significantly higher even in overweight/obese men than the lean individuals [63]. Adipokines, in particular leptin, mediate the crosstalk between adipose tissue and metabolic organs especially the liver, muscle, pancreas, and central nervous system, and plays a central role in appetite and body weight homeostasis[64], and *C. albicans* directly or indirectly inhibited excessive leptin and insulin release, although the mechanism involved requires further investigation. However, more ghrelin production helped in increased food intake and body weight gain in mice fed *C. albicans* mixed normal diet. In addition to insulin, the recent finding suggested the role of the resistin hormone associating obesity to type 2 diabetes and it represents a potential candidate for the etiology of insulin resistance[30]. As reported earlier, we found a higher level of resistin secretion in our DIO animal models indicating impaired glucose tolerance and insulin action that was mitigated by dietary *C. albicans*. Amylin is also co-secreted with insulin from the pancreatic β-cells, plays a role in glycemic regulation by slowing gastric emptying and promoting satiety, thereby preventing post-prandial spikes in blood glucose levels[65]. Surprisingly, although the blood RBS was unaltered in various groups of mice, amylin level was lowered in the diets containing *C. albicans*. Thus, the interaction of *C. albicans* with the host seems more complex than the simple concept of heredity and energy imbalance (**Fig. 10**). Nevertheless, dietary *C. albicans* has beneficial attributes in terms of metabolism for the host in both diets.

### Dietary *C. albicans* negates obesity induced pro-inflammatory immune activation

Several studies have shown that adults with overweight or obesity have a blunted lymphocyte response to mitogenic or antigenic stimulation, lower naïve lymphocyte output, a skewed T cell phenotype towards Th1 cells, and impaired macrophage and NK cell function. Numbers of certain types of lymphocytes, namely CD3^+^ cells, CD4^+^ and CD8^+^ T cell subsets, and B cells are also compromised in individuals with obesity[66, 67]. These deficiencies are thought to contribute to a higher risk of infections and chronic disease[68]. Leptin is known to regulate both innate and adaptive responses through modulation of immune cells activity, survival, and proliferation[64]. In innate immunity, leptin increases the cytotoxicity of natural killer (NK) cells and promotes the activation of granulocytes, macrophages, and DCs. In adaptive immunity, leptin increases the proliferation of naïve T cells and B cells while it reduces that of regulatory T cells (Treg). Leptin promotes the switch towards a pro-inflammatory Th1 (which secretes IFNγ) rather than anti-inflammatory Th2 (which secretes IL-4) phenotype and facilitates Th17 responses[64]. Accordingly, an increase in leptin seems to be associated with the decreased populations of macrophages, dendritic cells, WBC, lymphocyte, eosinophil, and lymphoid cells, and increased populations of myeloid cells and neutrophils in mice subjected to HFD than to a normal diet. However, the addition of *C. albicans* to the high-fat diet prevented this immune cells imbalance. We did not find the effect of diets on circulating T cells and granulocytes population, however dietary *C. albicans* somehow increased T cells but decreased granulocytes populations. Like T cells and macrophages, B cells also have been shown to play important roles in chronic inflammatory and autoimmune diseases, including systemic lupus erythematosus, rheumatoid arthritis, scleroderma, Type 1 diabetes, and multiple sclerosis[69], but only recently their role in obesity related insulin resistance have been revealed[70]. While the choice of diet did not affect B- cells and their subtypes except CD5^+^ B cells, the presence of *C. albicans* increased marginal zone and CD5^+^ B cells in a normal diet but reduced the population of follicular B cells. Fungal blended HFD only reduced these B cells populations. However, CD5^+^ B cells were more in the spleen of HFD diet fed mice than in mice with a normal diet indicating the development of autoimmune pathogenesis in obese mice. Thus, B cell manipulation could represent a novel approach to the treatment of obesity related insulin resistance and potentially to the prevention of T2D. In obese BALB/c mice, the spleen harbors an increased population of highly inflammatory TNFα^+^, IFNγTNFα double positive T helper cells along with low CTLA4 expressing CD4 T cell. Elevated levels of circulating TNFα have been linked to a wide variety of diseases, including arthritis, insulin resistance, Crohn’s disease, and cachexia associated with terminal cancer and AIDS[71]. IFNγ^+^ CD4 T cells that can stimulate NK cells and neutrophils are primary activators of macrophages.

Interestingly IFNγ^+^ CD4 T cells were found low in HFD than the mice fed with a non-obesogenic diet indicating a proinflammatory immuno-compromised state in obesity. IFNγTNFα double positive T helper cells in HFD while increasing IL17A producing CD4 T cells in both the diets fed mice suggesting Th17 mediated antifungal immunity possibly induced at the gut. However intra-peritoneal challenge of *C. albicans* in obese mice has shown a low Th17 mediated antifungal immunity with high CTLA4 expression CD4 T cells indicating obesity induced an immune-compromised state.

### Influence of dietary *C. albicans* on gut microbiota

Differential gut microbial composition in healthy *Vs* obese animals and humans has been demonstrated. Like in obese humans, this study also finds similar results where the proportion of Firmicutes and Bacteroidetes is associated with obesity[57]. Interestingly, despite pooling the fecal samples of four groups of mice, the metagenome analyses obtained here were very similar to the earlier reported individual mice or animal specific meta-data [13, 21, 58]. While the abundance of *Akkermansia* (phylum Verrucomicrobia) is negatively associated with body fat percentage growth in humans, relative abundances of *Lactoococcus* from phylum Firmicutes and with the genera *Allobaculum* (phylum Bacteroidetes) are positively associated with obesity[72]. We did not detect *Lactoococcus* and *Allobaculum,* rather *Parabacteroides distasonis* as a positively associated bacteria with obesity. Our study revealed enrichment of *Akkermansia muciniphila* in the gut of mice subjected to *C. albicans* mixed HFD only, suggesting its role in healthy body weight maintenance and energy expenditure. In addition to bacterial composition, gut mycobiota also changes in obese subjects. Human gut mycobiome harbors more than 66 genera and 184 species of fungi, while *Candida*, *Saccharomyces,* and *Cladosporium* predominate. Rodriguez et al. reported no significant differences in the richness of the mycobiome between non-obese and obese subjects, although the family biodiversity was significantly reduced in obese subjects[14]. Comparison to non-obese, obese subjects showed an abundance of fungal species from the class Saccharomycetes, families Dipodascaceae and Saccharomycetaceae, and a rare presence of fungi belonging to class Tremellomycetes. Fungal species from the genus Nakaseomyces and Pichia are also abundantly found in obese individuals. Although *Candida* sp. has been successfully identified from the intestine of healthy individuals and some studies have suggested a link between *Candida* sp. expansion and diabetes and inflammatory disorders, no differences in the relative presence of *Candida albicans* in obesity was observed[15, 16]. However, a recent study points out a positive correlation of *Candida albicans, Candida kefyr,* and *Rhodotorula mucilaginosa* fungal species with human obesity [17]. This study reports the first mycobiome analyses in animal models and we clearly demonstrated *C. albicans* enrichment in HFD as opposed to the normal diet fed mice. Since dietary *C. albicans* colonizes in the gut, one would expect its excessive abundance in fungal mixed HFD fed mice. However, while it synergized its own colonization in the mice feeding non-obesogenic food, interestingly, dietary *C. albicans* antagonized its growth in HFD fed mice. Additionally, it also altered other fungal abundance. The abundance of *Rhodotorula mucilaginosa*, *Trichosporon insectorum, Aspergillus ruber,* and *Rhizopus arrhizus* was negatively associated with obesity. Thus, diet form of *C. albicans* colonizes in the got and simultaneously it also modulates both bacterial and fungal ecosystems in the gut, and this modification could be one of the key regulatory factors responsible for the differential obesity parameters, immune cells variations, and immunity in different groups of mice used in this study.

Changes in microbial composition have been shown to alter host–microbiome interactions to mitigate diseases[73, 74]. Recent studies have evaluated the possibility of therapeutic targeting of the gut microbiome to reduce obesity by employing probiotics. For example, Stenman et al. targeted HFD induced dysbiosis by using *Bifidobacterium animalis* lactis 420 and found a decrease in fat deposits, improvement in glucose tolerance, and reduced inflammation[75]. Similarly, *Saccharomyces boulardii* Biocodex, a probiotic yeast, has been reported to reduce body weight, fat mass, and T2D in mice[76]. Mice have an endogenous level of *C. albicans* in the gut that increases substantially in HFD consumption or addition of it to the normal diet exogenously. However, dietary *C. albicans* somehow inhibits its excessive colonization when HFD is consumed. Thus, a threshold level of *C. albicans* colonization in the gut is needed to maintain eubiosis in the gut where microbes collectively facilitate the burning of excess calories in the high-fat diet, minimize abdominal fat storage to reduce obesity induced complications, and improve immunity (**Fig. 10**). Although HFD fed mice exhibited low immunity and abnormal immune cell profiling, dietary *C. albicans* did not cause any infections, rather it negated the HFD induced abnormalities. Therefore, we suggest a possible use of *C. albicans* as a probiotic especially in obese people or people dependent on high fat calorie intakes to manage obesity and associated complications. Since several genomic knockout strains of *C. albicans* were shown to be attenuated, such candidate strains may be the preferable probiotic isolates[60, 77]. In conclusion, this study is the first to show that *C. albicans* is not always an adversary rather a bonafide companion of the host.

## Materials and Methods

### Animals

All mouse experiments were conducted in strict accordance with the guidelines of the Institutional Animal Ethical Committee, Institute of Life Sciences, Bhubaneswar, India. Experimental mice protocols were fully approved by the committee and given the ethical Permit Number ILS/IAEC/133- AH/AUG-18. All efforts were made to minimize suffering and ensure the highest ethical and humane standards.

### Chemicals, reagents, and antibodies

High-fat diet (# 260HF, ∼36% fat) and Control diet (# D-131, ∼5% fat) were procured from SAFE® DIETS, Augy, France. Dulbeccòs Phosphate Buffered Salt Solution without Ca and Mg (# P04-36500), South American Fetal bovine serum (# P30-3302), and RPMI 1640 (# P04-16500) were purchased from PAN Biotech, GmbH, Germany. Phorbol 12-myristate 13-acetate (PMA), Ionomycin, Brefeldin-A, Albumin bovine serum- lyophilized powder, and Penicillin-Streptomycin (10,000 U/mL) were purchased from Sigma-Aldrich. LIVE/DEAD™ Fixable Blue Dead Cell Stain, and Dynabeads™ Untouched™ Mouse CD4 Cells Kit were purchased from Thermo Fisher Scientific Inc, MA, USA. YPD media (# G037-500g), Sulphosalicylic acid solution (3%), Benedict’s reagent were from HIMEDIA, India. EnzyChrom™ ketone body assay kit (EKBD-100) was from BioAssay Systems, Hayward, CA, USA. Dr. Morepen® GlucoOne, (Blood Glucose monitoring system Model BG-03) and BG- 03 test strips from Morepen Laboratories Ltd. Delhi, India. Periodic Acid Schiff (PAS) Stain Kit (Mucin Stain) (#ab150680) was obtained from Abcam, Cambridge, MA 02139-1517, USA. MILLIPLEX MAP Mouse Metabolic Hormone Magnetic Bead panel was purchased from MILLIPORE-Merck, Darmstadt, Germany, and used for the simultaneous quantification of Amylin (active), C-peptide 2, Ghrelin (active), GIP (total), GLP-1 (active), Glucagon, IL6, Insulin, Leptin, MCP1, PP, PYY (total), Resistin, and TNFα. BV510 Rat Anti-Mouse CD43 (Clone# S7), PE-CF594 Rat Anti-Mouse CD93 (Clone# AA4.1), and BD Cytofix/Cytoperm™ solution were purchased from BD Biosciences, San Jose, CA 95131, USA. Brilliant Violet 421™ anti-mouse CD21/CD35 (CR2/CR1) Antibody (clone#7E9), Alexa Fluor® 488 anti-mouse CD5 Antibody (clone#53-7.3), PE anti-mouse CD23 Antibody (clone#B3B4), Alexa Fluor® 700 anti-mouse Ly-6G/Ly-6C (Gr-1) Antibody (clone#RB6-8C5) and PerCP/Cyanine5.5 anti-mouse IL17A Antibody(clone# TC11-18H10.1) were purchased from BioLegend, Inc, San Diego, CA 92121, USA. PerCP-Cyanine5.5 Anti-Mouse CD19 (clone#1D3), APC Anti-Mouse NK1.1 (CD161) (clone#PK136), APC-Cyanine7 Anti-Human/Mouse CD11b (clone#M1/70), and Flow Cytometry Perm Buffer (10X) were purchased from TONBO biosciences, San Diego, CA, USA.

### Diet induced obesity and *C. albicans* in oral diet

The in-house-bred male BALB/c mice, about 4 to 5 weeks of age were co-housed and maintained under the standard conditions such as highly ventilated space with Hepa filtered air, maintained environment temperature, clean bedding, 12 hours to 12 hours dark-light cycle in individually ventilated cages. The initial body weight of each ear marked mouse was measured using a precision physical balance and further, it was measured after every 30 days. Age, body weight and blood glucose matched mice (4 mice per cage) were divided into different groups. The cages were supplemented with soft food dough (prepared with or without *C. albicans* in normal diet (ND) or high-fat diet (HFD)) and water provided *ad libitum* to the respective experimental groups for the duration of 21- 22 weeks as mentioned elsewhere. About 1 gm of freshly grown overnight cultured *C. albicans* cell pellet was mixed with 100 gm of powdered diet and the soft dough was prepared with sterile water. Similar consistency of food without fungal mix was also prepared. About 5 gm/mice food was provided daily morning and on an average ∼70-80 % of the food used to be consumed. Mice body weight and random blood glucose were monitored periodically until the sacrifice was due. Mice urine and fecal sample collection were carried out by placing the mice in metabolic cages 24-48 hrs prior to the sacrifice schedule on the 150th day of dietary intervention. On sacrifice day, under anesthesia (isoflurane inhalation) mice blood was collected via cardiac puncture using a 2 ml syringe and 26 gauge needle with potassium EDTA anticoagulants vials. Following immediate centrifugation at 4°C, plasma was separated and stored at -20°C until analysis. With cervical dislocation mice were sacrificed and dissected out for isolation of kidneys (tissue histology) and spleen (characterization of immunity).

### DIO Immune response to *C. albicans* intraperitoneal (IP) challenge

Normal diet (ND) or high-fat diet (HFD) DIO group of mice were supplemented with water and soft dough of ND or HFD ad libitum respectively for the duration of 21 weeks. At the end of dietary intervention, obese and control mice were challenged intraperitoneal (IP) with 6 x 10^5^ *C. albicans* cells in 100 μl of normal saline. 48 hours to post IP challenge, under anesthesia (isoflurane inhalation) mice blood was collected via cardiac puncture using a 2 ml syringe and 26 gauge needle with potassium EDTA anticoagulants vials. With cervical dislocation mice were sacrificed and dissected out for isolation of the spleen.

### Measurement of Blood Glucose

Under anesthesia, a tip of the mice’s tail was cut thereby a small drop of oozing out blood touched on GlucoOne BG-03 test strip in GlucoOne blood glucose monitoring system for the measurement of random blood glucose. The displayed values were noted and expressed the glucose as milligram per decilitre of blood.

### Estimation of Urine sugar

To a 1 ml of Benedict’s qualitative reagent, added with 100 μl of mouse urine mixed and boiled over a flame for 5-10 minutes, then cooled under the tap water, the content of the urine will be turned turbid due to precipitate that range from green to red depending on the increasing amounts of reducing sugar [78]. The formed turbidity of precipitate was measured at 650 nm using Eppendorf biophotometer plus.

### Urine protein estimation

Urinary total proteins were measured by precipitating the proteins with 3 % sulphosalicylic acid (SSA). The turbidity thus formed was compared with that of a protein standard bovine serum albumin at 650 nm. 1 part of urine or standard (1:2 serial diluted concentration with 10 mg/ml as maximum and 0.3125 mg/ml as minimum) were gently mixed with 4 parts of 3% SSA, and after 5 minutes at room temp (25-32°C) the absorbance of assay mixture was measured at 650 nm and 420 nm against reagent blank using VICTOR Nivo Multimode Microplate Reader or Eppendorf biophotometer plus in different experiments. A known concentration of protein solution was used to prepare a standard curve in the range from 0 to 10 mg/ml. The unknown urinary protein concentration was then determined from a plot of concentration versus absorbance obtained for the standard protein solutions [79].

### Histopathology

Mouse kidney paraffin sections were stained with Periodic Acid Schiff reagents and counterstained with Mayer’s Hematoxylin as per the manufacturer’s instructions [60].

### Ketone body analysis

Concentrations of Acetoacetic acid (AcoAc) and 3- hydroxybutyric acid (BOH) were measured in plasma or urine samples from mice using EnzyChrom ketone body assay kit as per the manufacturer’s instructions. The assay is based on 3-hydroxybutyrate dehydrogenase catalyzed reactions, in which the change in NADH absorbance, measured at 340nm, is directly related to the AcoAc and BOH concentrations. At pH=7.0, AcAc + NADH + H+ ⇋ BOH + NAD^+^; the reverse reaction happens at pH=9.5, AcAc + NADH + H+ ⇋ BOH + NAD^+^. As per the assay guide a concentration of *AcoAc* = (*ODblank* –*OD sample*) ÷ (*ODH*20 −*ODstandard*) × 8 and *BOH*= (*ODsample*–*OD blank*) ÷ (*ODstandard* –*ODH*20) × 8, and were expressed in millimolar (mM) concentration of respective ketones. For the calculation of total ketone body (TKB) concentration, *TKB* = (*AcoAc* +*BOH*) and expressed in mM.

### Mouse hormone Assay

Leptin, Ghrelin, Resistin, Amylin, Gastric inhibitory polypeptide (GIP), Glucogon-like peptide-1 (GLP-1), Pancreatic polypeptide (PP), Insulin, Glucogon, C-peptide, and Peptide tyrosine tyrosine (PYY) were measured from plasma using mouse multiplex magnetic bead based metabolic hormone panel as per manufacturers protocol. The samples, standard, control, and blanks in the panel were acquired in Bio-plex 200 system and analyzed using 5PL algorithms in Bio-plex Manager 5.0 software.

### Absolute Blood cell counts

Mouse blood was collected in Potassium EDTA vials mixed well, about 50 μl of blood sample was taken in a capillary tube specifically for automated blood cell counter (Exigo Eos Veterinary Hematology Analyser, Sweden) and an absolute value of each cell was estimated.

### CD4^+^ T cell Isolation

Untouched CD4^+^ T cells (negative separation) were isolated from mouse spleen with magnetic Dynabeads kit as per the manufacturers’ instructions. Briefly, splenocytes were prepared by mashing up the mice spleen in a 40 µm sieve cell strainer with the plunger end of the syringe in a petri dish containing RPMI-1640 complete medium (10 % Fetal Bovine Serum (FBS), Penicillin and streptomycin supplemented). Spleenocytes were washed in PBS (without Ca2^+^ and Mg2^+^) containing 0.1% BSA and 2 mM EDTA, pH 7.4, and adjusted the cell concentration into 1 x 10^7^ cells/ml. Added a cocktail of antibodies that binds non CD4^+^ T cells, followed by addition streptavidin secondary antibodies magnetic dynal beads depletes all cells except CD4^+^ T cells. Thus, collected CD4^+^ T cells were washed in complete medium or 0.1% BSA-PBS, and used for the subsequent experimental protocol.

### Multicolour Immune phenotyping of tissue resident and peripheral immune cells

Splenocytes or blood cells (50-100 μl of whole blood) was washed in PBS (without Ca2^+^ and Mg2^+^) containing 0.1% BSA, pH 7.4 and RBC lysed in ACK lysing buffer following stop lysis using 2 % FBS, 2 mM EDTA in phosphate buffered saline pH 7.4 and centrifuged at 380x g for 8 min. The cell pellet again washed lysing buffer. Staining was carried out with the following fluorescent tagged antibodies such as APC-Cy7 anti- mouse CD11b, Alexa flour 488 anti-mouse CD5, PerCP-Cy5.5 anti- mouse CD19, Brilliant violet 421 anti- mouse CD21/35, PE anti- mouse CD23, Alexa flour 700 anti- mouse Gr-1 (Ly6G/Ly6C), Brilliant violet 510 anti- mouse CD43, and Live dead blue fixable dye. After washing twice, the stained cells were acquired in BD LSRFortessa and analyzed using FACSDIVA software ver 8.2.

### Characterization of CD4^+^T cell response

Untouched CD4^+^ T cells were activated with 50 ng/ml PMA and 1 μg/ml Ionomycin for 4 hours under standard aseptic cell culture condition in RPMI 1640 complete medium, on the 3rd hour 10 μg/ml of brefeldin A was added. Finally, the cells were harvested and washed twice in 2% FBS, 2mM EDTA in phosphate buffered saline pH 7.4 and stained with APC anti-mouse CTLA4, followed by live dead blue staining, washed further and fixed the cells in BD Cytofix/Cytoperm, fixation/permeabilization solution. The fixed cells were washed and permeabilized in 500 μl of BD Perm/Wash Buffer, stained with the fluorescently tagged antibodies such as Alexa flour 488 anti-mouse Interferon-gamma; Alexa flour 700 anti-mouse tumor necrosis factor-alpha; and PerCPcy5.5 Interleukin- 17A antibodies. After washing twice, the stained cells were acquired in BD LSRFortessa or Cytoflex S and analyzed using FACSDIVA software ver 8.2 or CytExpert software respectively.

### Microbial DNA extraction from mice fecal sample

The microbial metagenomic DNA from the mice faecal samples were extracted by using a well-established Guanidinium thiocyanate (GITC) based protocol with a minor modification[80]. Briefly, about 200 mg of stool sample was resuspended in Tris–EDTA buffer (pH 8.0), homogenized using sterile glass beads (2.5 mm), and the suspension was collected after centrifugation followed by enzymatic lysis using a cocktail of lysozyme (10 mg/ml) (MFCD00131557 - Mp Biomedicals India Pvt. Ltd), Zymolyase (20 mg/ml) (08320921- Mp Biomedicals India Pvt. Ltd) and lyticase (20 mg/ml) (37340-57-1 -Sigma-Aldrich) at 37 °C overnight. The next day, 250 µl of 4 M GITC was added to the suspension, vortexed for 1 min, then 120 µl of 10 % N-Lauryl sarcosine and 650 µl 5% N-Lauryl sarcosine were added, respectively with an interval of 10 min. The mixture was vortexed and incubated at 70 °C for 1 hour. The mechanical cellular disruption was carried out with 0.1 mm silica beads (1177 P71-Mp Biomedicals India Pvt. Ltd)) in FastPrep 24 5G homogenizer (Mp Biomedicals India Pvt.

Ltd) 3 times with a cycle of 30s beating and 30s rest, and followed by washing in 1 ml of Polyvinyl Polypyrollidone (PVPP)(77627-Sigma-Aldrich). The supernatant was collected from the homogenized samples by centrifugation, mixed gently with 2 ml isopropanol, and incubated overnight at 4 °C. After incubation, crude DNA was precipitated by centrifugation and dried at 42 °C for 1 hour. The dried pellet was resuspended in 900 µl of PBS and 100 µl of 0.5M Potassium acetate and incubated at -20 °C for 1 hour, followed by incubation at 4 °C for 30 min. RNase A (10 mg/ml) (DS0003-Himeda) treatment was carried out for 30 min at 37 °C. Finally, DNA was extracted by using 96 % chilled ethanol and spinning at 14,000 rpm for 30 min at 4 °C. The pellet was dried at 42 °C for 1 hour and suspended in 60-100 μl of TE buffer (pH -8.0) depending upon the pellet amount. The DNA concentration was determined in NanoDrop spectrophotometer, Qubit dsDNA HS Assay Kit (Q32854, Invitrogen, USA) and further verified in Agarose gel electrophoresis.

### Semi-quantitative PCR for *C. albicans* detection in fecal DNA

Isolated total DNA from the fecal samples of the four mice groups were labelled as BND, BCND (Balb/c mice subjected to a normal diet without or with *C. albicans* mix, respectively), BHFD, and BCHFD (Balb/c mice subjected to a high-fat diet without or with *C. albicans* mix, respectively). To identify the presence of *C. albicans* DNA in these samples, and semi-quantitative end point PCR was performed to amplify CaPCNA and CaEfg1. PCR reaction was carried out in a 100 µl master mix having dNTPs (200 µM), 20 µl 1X Taq-DNA polymerase buffer, 2 Units of Taq-DNA polymerase, 20 picomoles of CaPCNA (NAP31 5’- GGC CAA GCT TGG ATC CAC ATA TGT TAG AAG GTA AAT TTG AAG-3’; NAP32 5’- GGC CGA ATT CGG ATC CCT ACT CAT CAT CAT CG-3’) OR CAEFG1 (NAP393 5’- CCG GGG TAC CGG ATC CAC ATA TGT CAA CGT ATT CTA TAC; NAP395 5’- GGC CCT CGA GAC CAC TAG GAG CAC TTG TG-3’) primers and 200 ng of total DNA as a template. The PCR cycling condition includes initial denaturation of 1 min. at 95 °C followed by 25 cycles of 95°C for 30s, 52°C for 30s,72 °C for 30s with a final extension of 3 min. The PCR products were separated on 1 % agarose gel and the image was taken in Bio-Rad ChemiDoc MP Imagers System.

### CFU Analysis

Fungal colonisation was estimated in the fecal samples by CFU determination. About of 200 mg of the samples from various groups were separately homogenised in sterile PBS and diluted to various concentrations and further spreaded onto YPD agar containing chloramphenicol (25 mg ml−1) to count for CFU. Random colonies were picked and colony PCR was conducted to confirm presence of *C. albicans* species. PCR amplification was carried out for CaPCNA gene.

### Metagenomics analyses

The composition of bacterial and fungal species in the gut of various groups of mice was determined by 16S and ITS metagenomics analyses of fecal DNA samples. After the quality control pass libraries were constructed in alignment with the 16S metagenomic library preparation protocol for bacteria and archaea from Illumina Inc. Briefly, 12.5 ng DNA was subjected to 16S V3-V4 PCR using the gene-specific primers (Fp 5’-TCG TCG GCA GCG TCA GAT GTG TAT AAG AGA CAG CCT ACG GGN GGC WGC AG-3’ and Rp 5’-GTC TCG TGG GCT CGG AGA TGT GTA TAA GAG ACA GGA CTA CHV GGG TAT CTA ATC C-3’) that target the V3 and V4 region of 16S rDNA[81]. Similarly, for the ITS analyses, the Fungal Metagenomic Sequencing Demonstrated Protocol from Illumina was used. The gene-specific sequences used in this protocol target the fungal ITS1 region between the 18S and 5.8S rRNA genes [82]. Briefly, 12.5 ng DNA was subjected to PCR using ITS1F/1R (5’–TCG TCG GCA GCG TCA GAT GTG TAT AAG AGA CAG CTT GGT CAT TTA GAG GAA GTA A- 3’ and 5’–GTC TCG TGG GCT CGG AGA TGT GTA TAA GAG ACA GGC TGC GTT CTT CAT CGA TGC-3’) and ITS4F/4R (5’–TCG TCG GCA GCG TCA GAT GTG TAT AAG AGA CAG CCC GGT CAT TTA GAG GAA GTA A-3’ and 5’ –GTC TCG TGG GCT CGG AGA TGT GTA TAA GAG ACA GGC TGC GTT CTT CAT CGA TGT-3’) primers.

Both the PCR products from all the four cohorts were bead purified and subjected to another round of PCR with dual indices and adapters to generate the respective libraries as recommended by Illumina. The cleaned libraries were quantitated on Qubit® flurometer and appropriate dilutions loaded on a D1000 screen tape to determine the size range of the fragments and the average library size. Libraries were diluted to 4 nM, pooled, spiked with 20 % PhiX pre-made library from Illumina, and loaded on a MiSeq v3 kit. Sequencing was performed for 2X300 cycles. The original raw data obtained from Illumina Miseq were recorded in FASTQ files. The reads obtained from the instrument were adapter free. The paired-end reads were assembled, filtered, trimmed, and aligned to SILVA 16S database using Mothur software v.1.44.1. The reads were clustered based on similarity and further clustered into OTUs (Operational Taxonomic Unit) at 97 % identity against greengenes database “gg_13_8_99”. For ITS analyses, Qiime2 q2cli version

2021.2.0 was used to analyze the raw reads after removing the low quality, noisy, unpaired, and chimeric reads by DADA2. The 118 amplicon sequence variants (ASVs) from the analysis were classified using UNITE dynamic classifier. The relative abundances were calculated using R. Relative abundance of top 10-20 phylum, class, order, family, genus, and species were determined. Taxonomic lineages at rank 11 or higher w.r.t. relative abundance, were collapsed as “Others”. Relative abundance (in percent) of each lineage is tabulated at each level. Unidentified categories are the ones for which taxonomy is not presently available. It’s different from unclassified, which means the reads could not be classified by the classifier.

### Statistical Analysis

Statistical Analysis of data sets derived from flow cytometric analysis, multiplex hormone panel, and quantification parameters were carried out using GraphPad Prism 8.0 and Microsoft Excel. Statistical significance was calculated based on two-way or one- way ANOVA, with Tukey’s post hoc multiple comparison test. Correlation and Regression analysis was carried out using Person’s correlation tests using GraphPad Prism 8.0.

## Supporting information

All the supplementary Informations

## Availability of data and materials

Metagenomics for bacterial and fungal DNA sequencing data have been uploaded to NCBI as PRJNA765929.

## Authors’ contributions

NA and DP conceptualized, designed, and analysed the study; DP, SRS, PK, and BU conducted experiments, analysed the data, and prepared initial draft of the manuscript. NA and DP wrote the final draft. NA obtained the funding. All the authors reviewed the results and approved the final version of the manuscript.

## Acknowledgements

We thank Mr. Sitendra Prasad Panda and Mr. Paritosh Nath for their technical assistance and our laboratory colleagues for their thoughtful discussion. Mr. Biswajit Patro for his technical assistance and nurturing of the animals was highly appreciated. For metagenomics, miBiome therapeutics is highly acknowledged. We thank DP for DBT-RA, and SRS and BU for CSIR-Senior Research Fellowships.

