## Supplementary material for "Commensal fungi *Candida albicans* modulates dietary high-fat induced alterations in metabolism, immunity, and gut microbiota": All the supplementary Informations

***Correspondence to:**

**Supplementary Figure 1: Effect of dietary C. albicans on DOI induced body weight and hormones (A)** A kinetics of body weight gain with respect to duration from mice fed with or without *C. albicans* in normal diet (ND) or high fat diet (HFD). Mean and standard error mean of various metabolic hormones from each of the groups of mice blood on the 150th day (~22 weeks) of dietary intervention, (**B**) plasma leptin to ghrelin ratio, (**C**) peptide tyrosine tyrosine (PYY) (pg/mL), (**D**) Amylin (pg/mL), and (**E**) Glucagon level (pg/mL). A linear regression and Person's correlation analysis of Insulin level versus leptin level from individual mouse irrespective of dietary intervention (**F**). A pie chart depicting different percentage of various metabolic hormones measured from various groups of mice (**G)**. A statistical significance (*p≤0.05, ** p≤0.01,***p≤0.001, **** p≤0.0001) was calculated using one-way ANOVA and Tukey's multiple comparison test.

**Supplementary Figure 2: Structure and function of kidney upon dietary manipulation**. (**A**) A representative image with table depicts urine glucose level by a semi quantitative Benedict's test. (**B**) Mean and standard error of mean of a kidney weight (gram) from all individual mice from the 4 groups of mice. (**C**) A representative images of kidney longitudinal section (2 micron thickness) from all 4 groups of mice were stained with periodic acid schiff's, counter stained with mayer's hematoxylin and analyzed using brightfield 40x objective, ZEISS ApoTome Microscope. A statistical significance (*p≤0.05, ** p≤0.01, ***p≤0.001, **** p≤0.0001) was calculated using one-way ANOVA and Tukey's multiple comparison test.

**Supplementary Figure 3: Tissue resident myeloid and lymphoid immune cells.** (**I**) A representative bivariate density plot of CD11b (y-axis) vs Side scatter (x-axis) for the analysis of percent total Myeloid and Lymphoid population in spleen using BD FACSDiva Software v8.0.2. Mean and standard error of mean of percent total tissue resident **(II)** Myeloid cells, and (**III**) Lymphoid cells from spleen of all the 4 groups mice on the of sacrifice, and a statistical significance (*p≤0.05, ** p≤0.01,***p≤0.001, **** p≤0.0001) was calculated using one-way ANOVA and Tukey's multiple comparison test.

**Supplementary Figure 4: Metagenomics analyses.** Various workflow followed in the metagenomic analyses of fecal samples (**A**) 16S rDNA and (**B**) 18S rDNA ITS1 analyses

**Supplementary Figure 5: 16s rDNA sequence analyses. (A)** Alpha rarefaction plot generated using Simpson to measure average bacterial species diversity within a sample. Taxonomic classification of OTUs was carried out and assigned them into top 20 classes (**B**), top 20 orders (**C**) and top 20 families (**D**). BND, metagenomic DNA isolated from normal diet fed BALB/c mice fecal sample; BCND, metagenomic DNA isolated from normal diet with *C. albicans* mix fed BALB/c mice fecal sample; BHFD, metagenomic DNA isolated from high fat diet fed BALB/c mice fecal sample; and BCHFD, metagenomic DNA isolated from normal diet with *C. albicans* mix fed BALB/c mice groups fecal sample.

**Supplementary Figure 6: ITS sequence analyses. (A)** Alpha rarefaction plot generated using Simpson to measure average fungal species diversity within a sample. Taxonomic classification of OTUs was carried out and assigned them into top 20 classes (**B**), top 20 orders (**C**) and top 20 families (**D**). BND, metagenomic DNA isolated from normal diet fed BALB/c mice fecal sample; BCND, metagenomic DNA isolated from normal diet with *C. albicans* mix fed BALB/c mice fecal sample; BHFD, metagenomic DNA isolated from high fat diet fed BALB/c mice fecal sample; and BCHFD, metagenomic DNA isolated from normal diet with *C. albicans* mix fed BALB/c mice groups fecal sample.

**Supplementary Figure 7: Overview of differential effect of dietary *C. albicans* on mice**. A brief summary of the alternation in different metabolic parameters, immune cells profiles, immunity and microbiome in mice subjected different calorie content diets without and with *C. albicans* probiotic**.** Colour coding has been inserted in a box.

**Supplementary Table 1: Kinetics of body weight gain in BALB/c mice.** Individual mouse were ear marked and tracked for change in body weight as per the mentioned duration upon diet challenge and presence or absence of *C. albicans* mix.

**Supplementary Table 2: Validity of metagenomics.** Read summary obtained from Illumina sequencing for each sample.

**Supplementary Table 3: Diversity and abundance of bacteria and fungi in the mice gut.** Percent abundance of top 20 bacterial and fungal species.

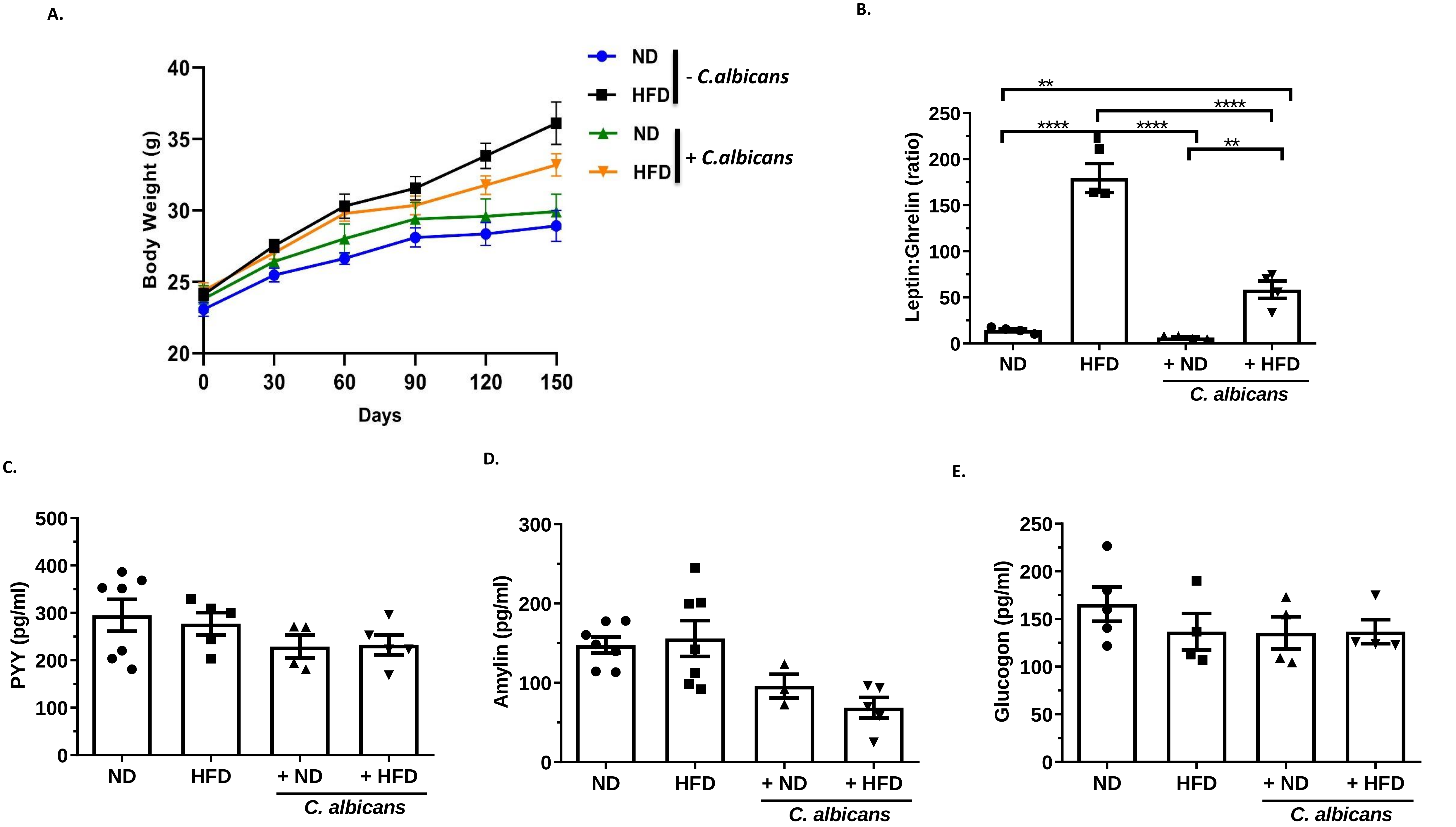

Supplementary Fig.1

F.

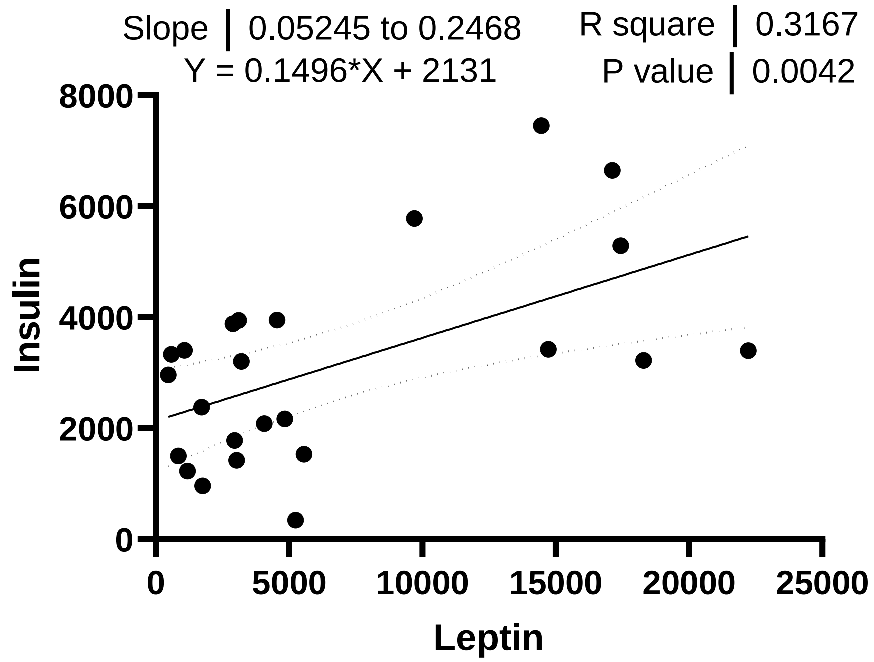

G.

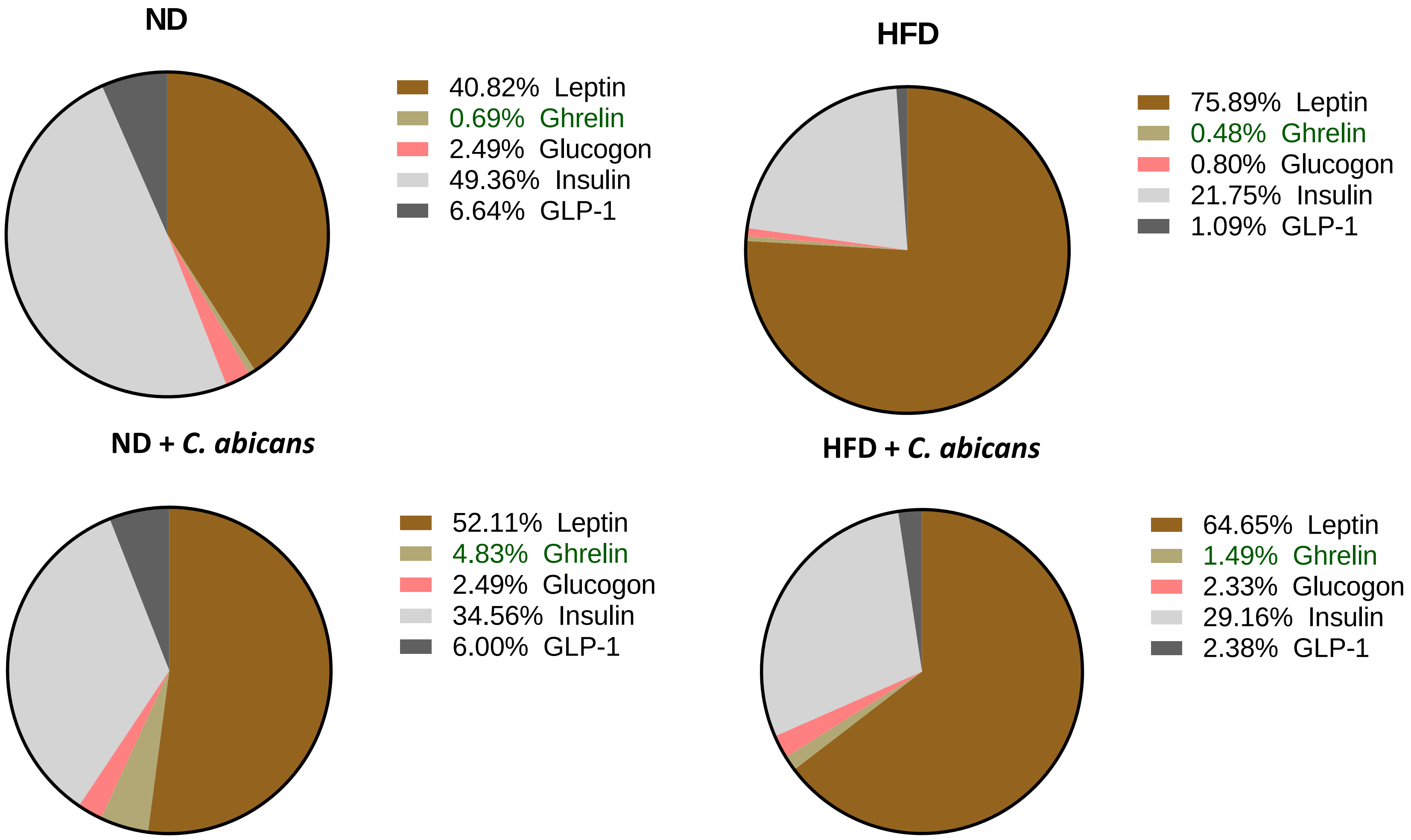

Supplementary Fig.1

A.

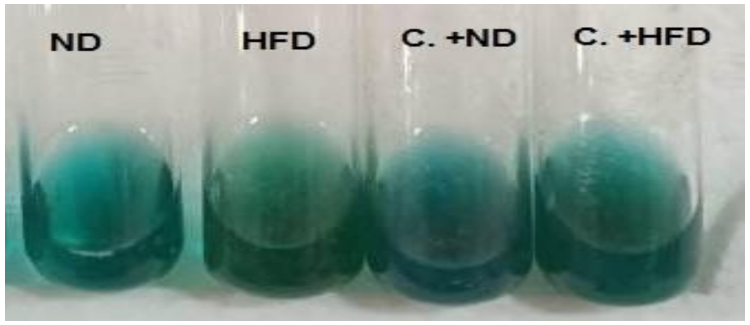

| Urine of Diet Conditions | Benedict's Semi quantitative read out |
|---|---|
| ND | nil ( No color change) |
| HFD | + (Green) |
| ND+ C. albicans | Trace/nil (Bluish Green) |
| HFD+ C.albicans | Trace/nil (Bluish Green) |

B.

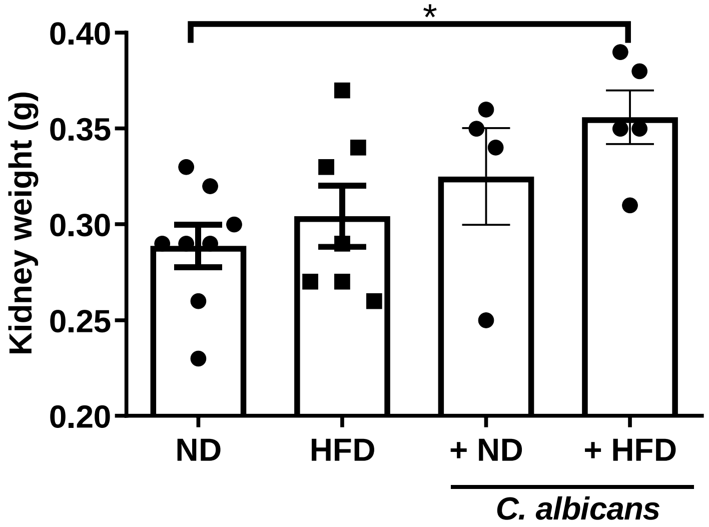

C.

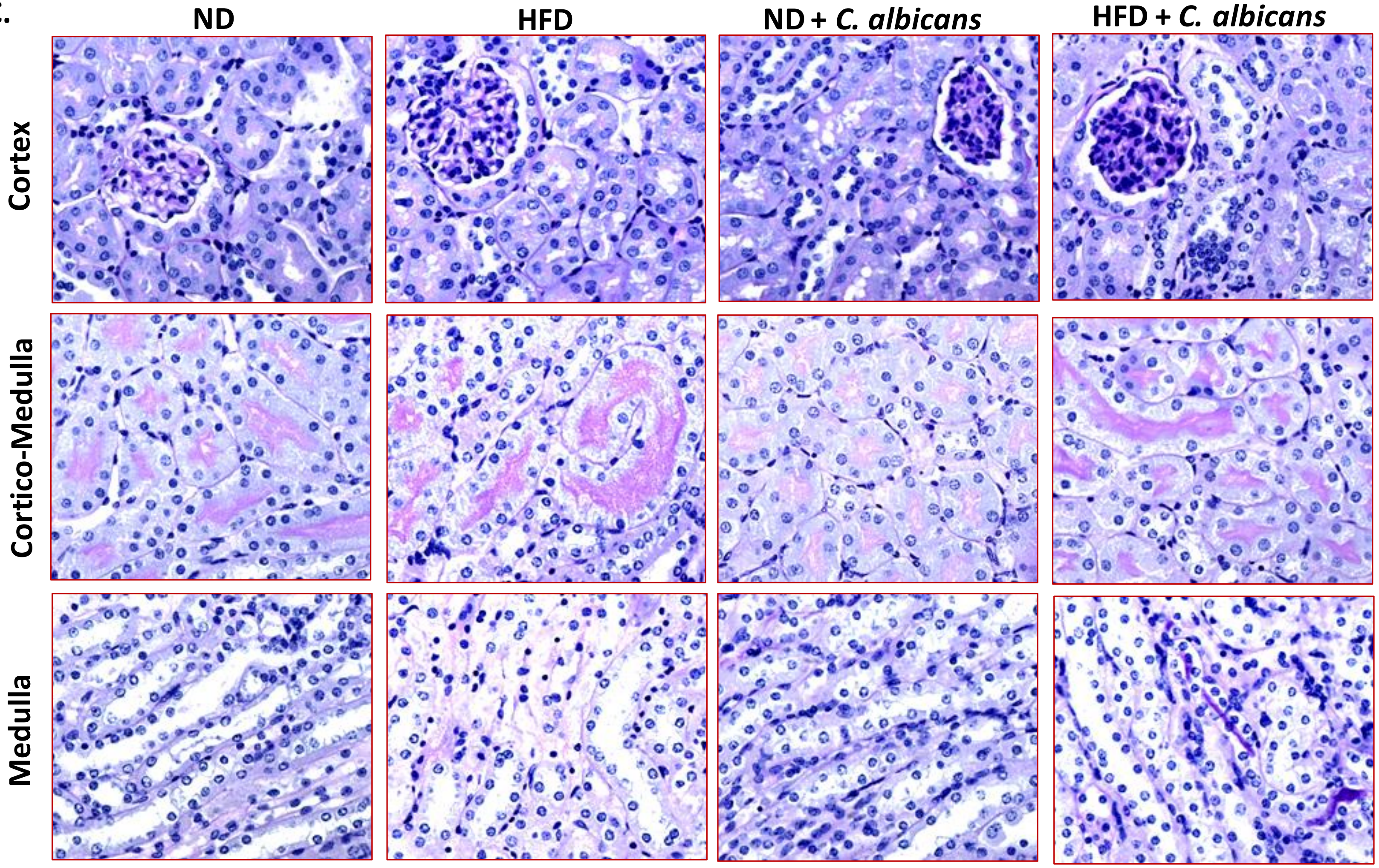

Supplementary Fig. 2

Tissue resident myeloid and lymphoid compartment

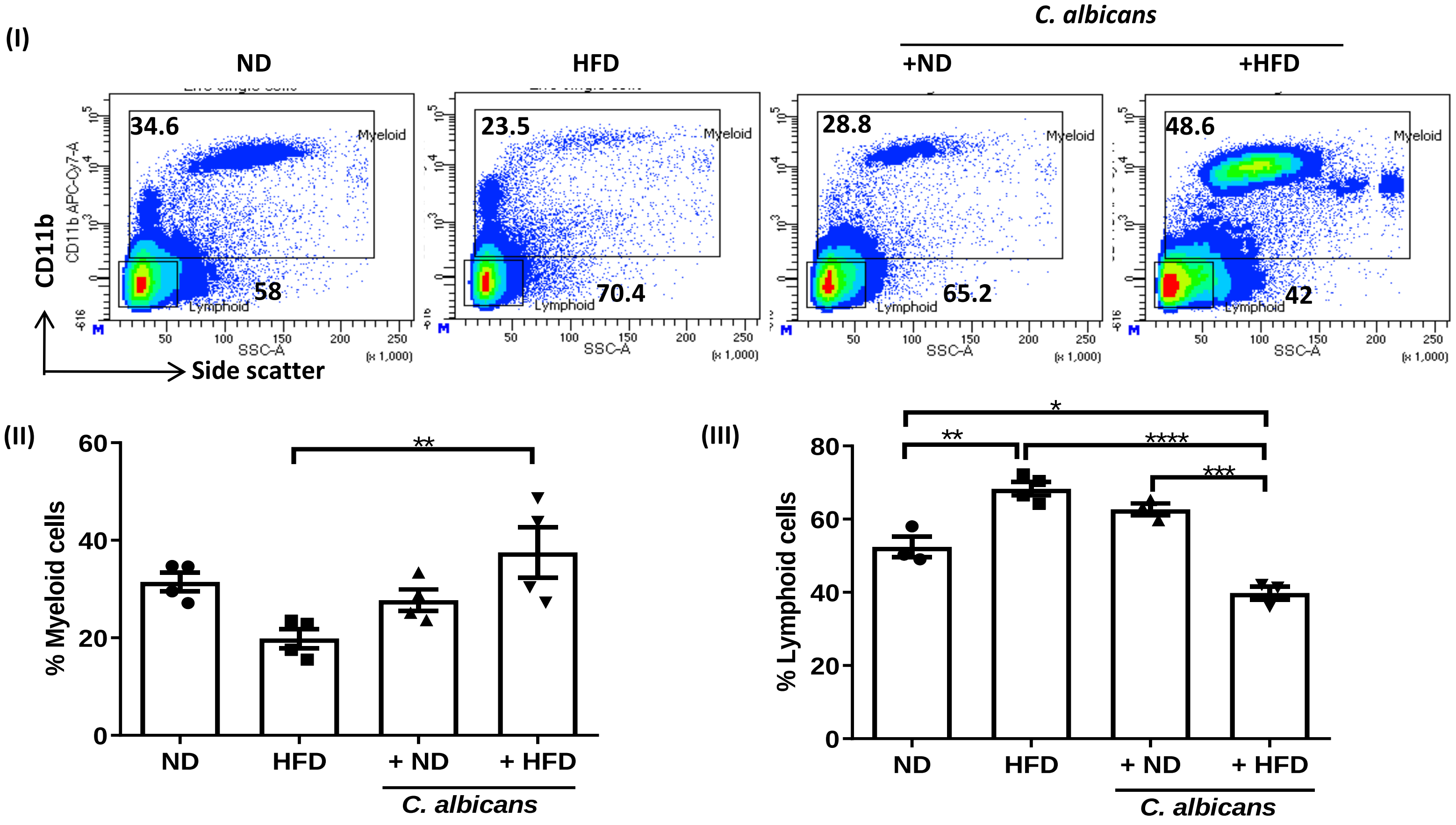

Supplementary Fig.3

#### A. Workflow for 16S rDNA Analyses

#### B. Workflow for ITS Analyses

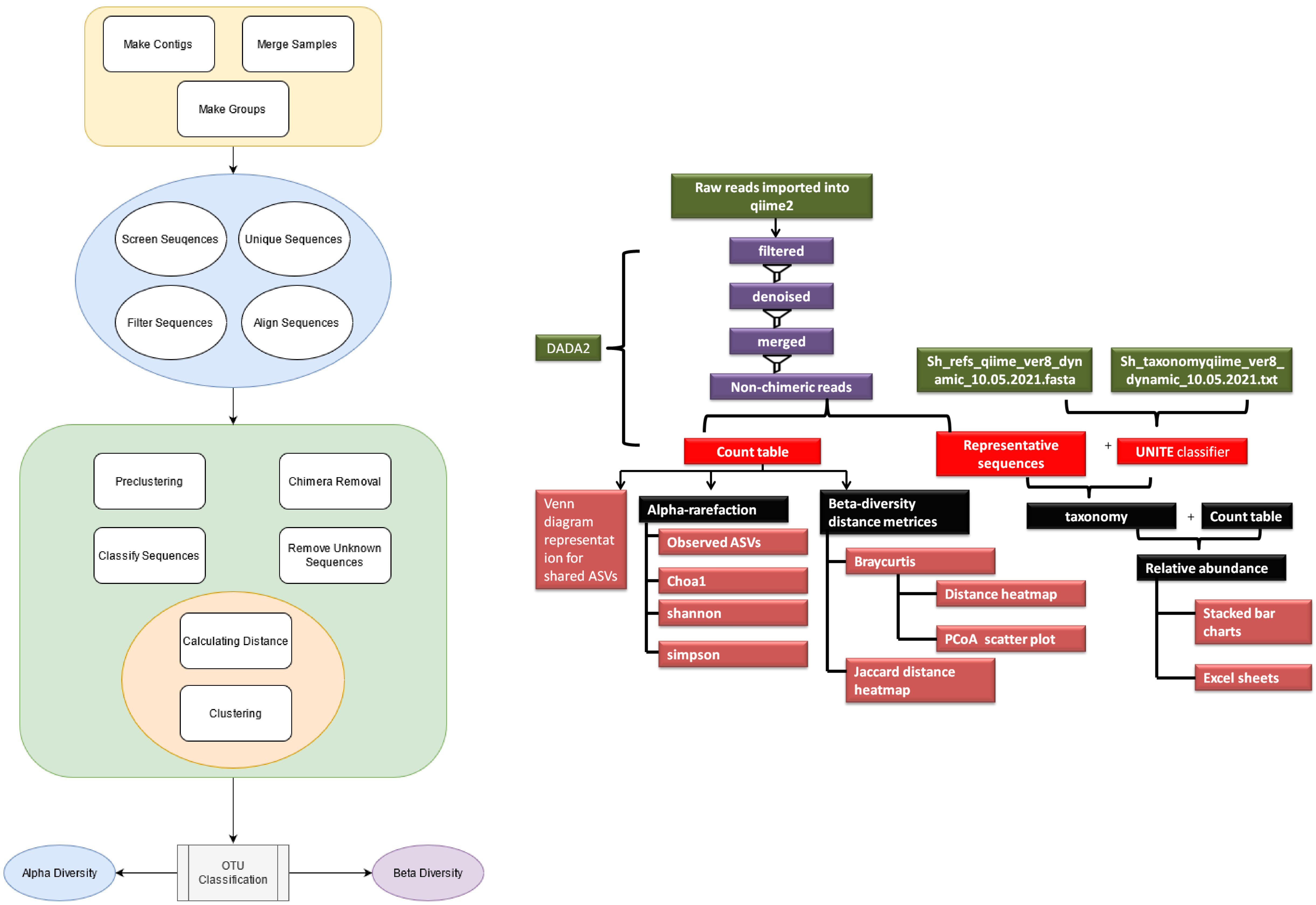

Supplementary Fig. 4

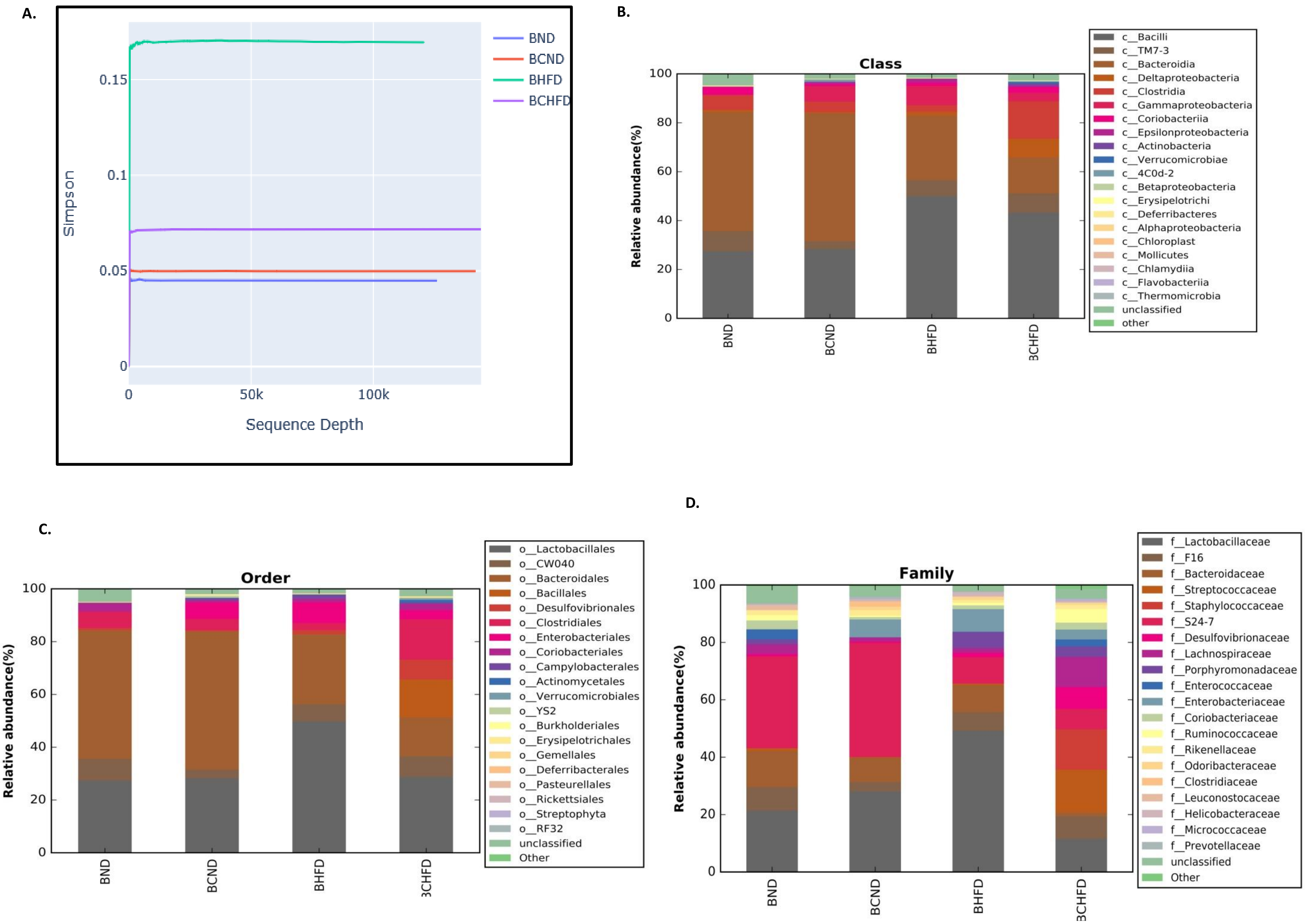

**Suppl. Figure 5**

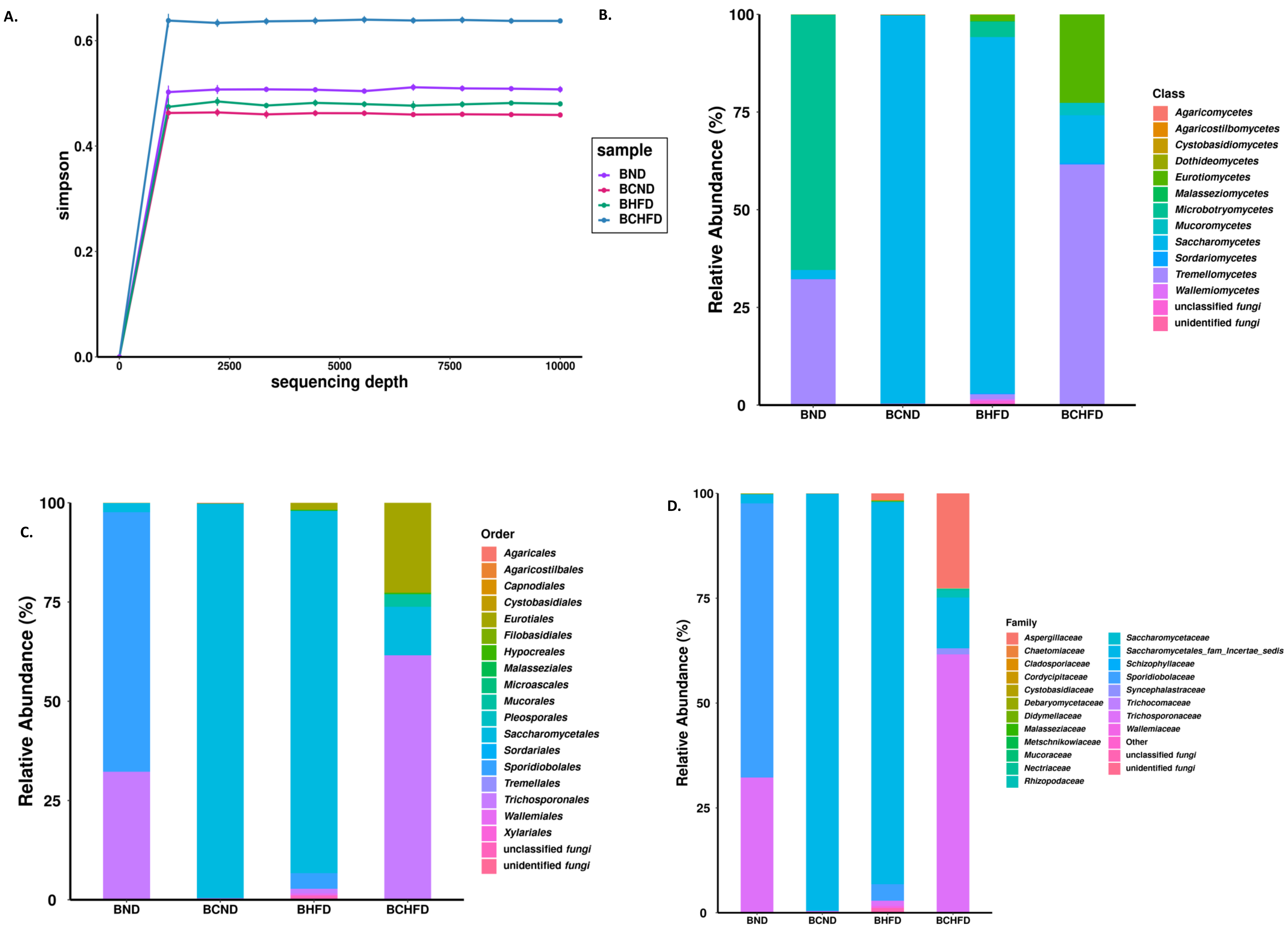

**Suppl. Figure 6**

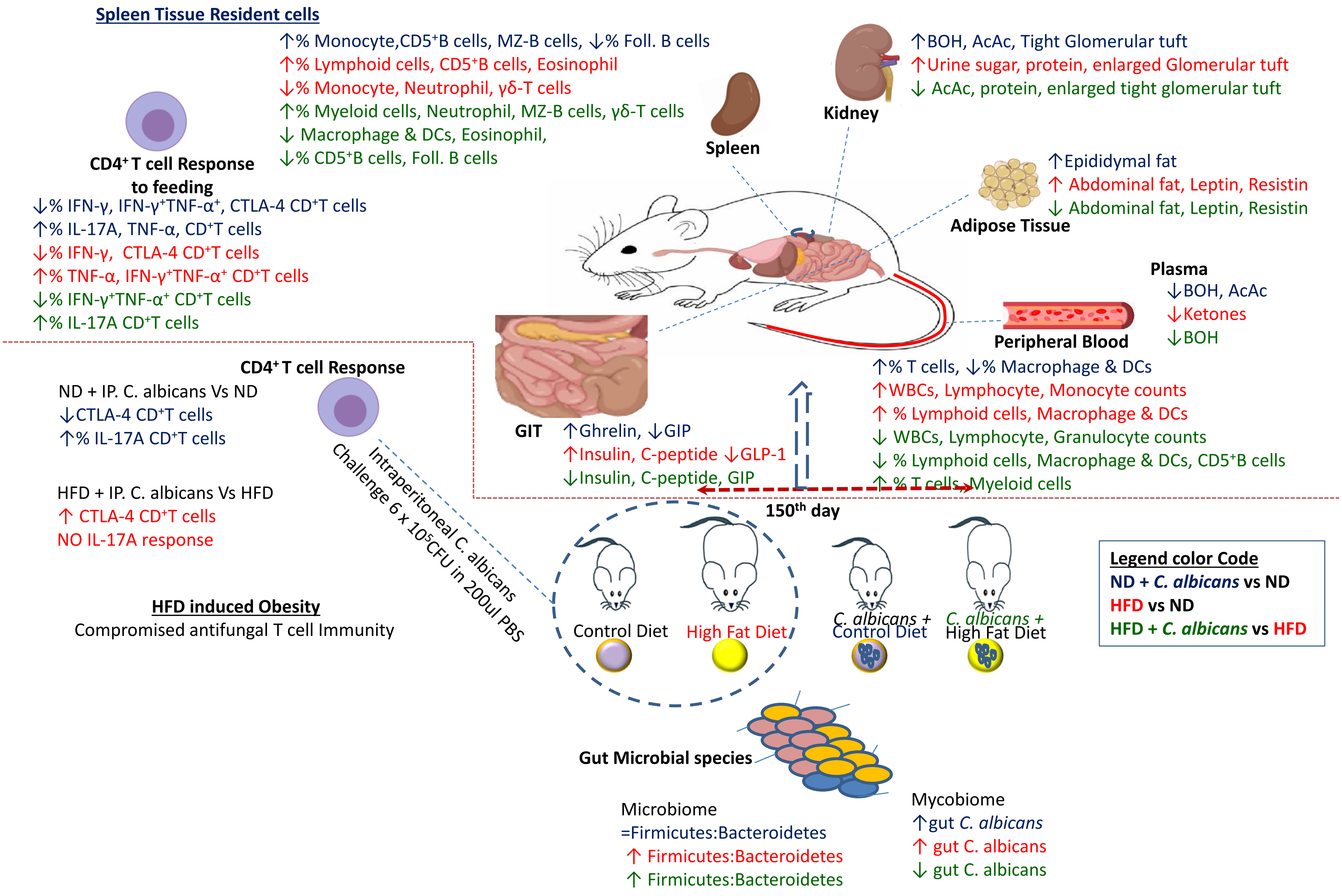

Supplementary Figure 7

**Supplementary Table 1:** Individual mouse were ear marked and tracked for change in body weight

| Body Wt(g) | Normal Diet | | | | | | High Fat Diet | | | | | |
|---|---|---|---|---|---|---|---|---|---|---|---|---|
| **DAYS** | 0 | 30 | 60 | 90 | 120 | 150 | 0 | 30 | 60 | 90 | 120 | 150 |
| **Without *Candida albicans* in diet** | 25.1 | 27.8 | 29 | 32.1 | 33.4 | 36.2 | 24.2 | 28.6 | 30.7 | 30.6 | 34.1 | 35.9 |
| | 24.9 | 27.2 | 27.6 | 29.6 | 29.7 | 29.5 | 24.6 | 26.6 | 26.8 | 27.4 | 29.3 | 31.1 |
| | 23 | 25.3 | 26 | 26.7 | 26.7 | 26.9 | 23.6 | 28 | 34.5 | 34.6 | 38 | 45 |
| | 22.5 | 25.5 | 26.4 | 27 | 27.2 | 27.9 | 25.8 | 27.1 | 32.4 | 33.9 | 33.6 | 33.8 |
| | 21.2 | 24.2 | 25 | 26.9 | 27 | 27.3 | 24.2 | 26.9 | 28.2 | 31 | 31.8 | 32.6 |
| | 22.1 | 24.2 | 26.1 | 27.3 | 27.5 | 27.5 | 24.4 | 26.6 | 30.8 | 32.7 | 35.1 | 36.4 |
| | 23 | 25 | 26.4 | 26.9 | 26.7 | 27.1 | 24.6 | 30 | 30 | 32.4 | 34.6 | 37.1 |
| | 22.8 | 24.6 | 26.7 | 28.3 | 28.7 | 29 | 21.5 | 26.4 | 29.1 | 29.9 | 34.1 | 37 |
| **Mean** | 23.1 | 25.5 | 26.6 | 28.1 | 28.4 | 28.9 | 24.1 | 27.5 | 30.3 | 31.6 | 33.8 | 36.1 |
| **SEM** | 0.5 | 0.5 | 0.4 | 0.7 | 0.8 | 1.1 | 0.4 | 0.4 | 0.9 | 0.8 | 0.9 | 1.5 |
| **Weight gain (%)** | 0 | 10.39 | 15.15 | 21.64 | 23 | 25.1 | 0 | 14.1 | 25.72 | 31.1 | 40.2 | 49.8 |
| **With *Candida albicans* in diet** | 22.4 | 24.7 | 26.3 | 27.2 | 27.3 | 27 | 23.4 | 27.1 | 27.9 | 29 | 30.2 | 30.4 |
| | 25.7 | 29.1 | 29 | 30.3 | 30.1 | 29.6 | 22.2 | 24.7 | 29 | 29.5 | 29.8 | 31.5 |
| | 22 | 24.6 | 26 | 27.3 | 27.4 | 28 | 23.3 | 26.7 | 29.9 | 28.2 | 31.3 | 32.3 |
| | 19.6 | 22.3 | 24.1 | 25.8 | 25.9 | 26.2 | 23.3 | 26.6 | 28.7 | 30.6 | 33.6 | 36.4 |
| | 26.4 | 29.5 | 32 | 34.8 | 35.4 | 35.2 | 24.5 | 28.2 | 30.8 | 31.4 | 32.3 | 34.3 |
| | 25.5 | 27.1 | 30.3 | 31.4 | 31.7 | 32.8 | 26.1 | 28.3 | 31 | 32.9 | 33.3 | 34.9 |
| | 25 | 27.7 | 28.5 | 29.1 | 29.3 | 30.7 | 26.5 | 28.3 | 32.4 | 32.8 | 34.2 | 35 |
| | 22.3 | dead | dead | dead | dead | dead | 25.8 | 26.4 | 28.6 | 28.5 | 29.5 | 30.7 |
| **Mean** | 23.8 | 26.4 | 28 | 29.4 | 29.6 | 29.9 | 24.4 | 27 | 29.8 | 30.4 | 31.8 | 33.1 |
| **SEM** | 0.9 | 1 | 1 | 1.2 | 1.2 | 1.2 | 0.6 | 0.4 | 0.5 | 0.7 | 0.6 | 1.2 |
| **Weight gain (%)** | 0 | 10.9 | 17.64 | 23.5 | 24.4 | 25.6 | 0 | 10.6 | 22.1 | 24.6 | 30.3 | 35.6 |

#### Supplementary Table 2

| Samples | Read length (bp) | Raw reads | Total reads in bp | Reads after filtration | #OTU /#ASV | Alpha Diversity Indices (Simpson) |
|---|---|---|---|---|---|---|
| **16S RNA Analyses** | | | | | | |
| **BND** | 35-301 | 272568 | 163540800 | 126118 | 1500 | 0.044 |
| **BCND** | 35-301 | 324926 | 194955600 | 141864 | 1628 | 0.049 |
| **BHFD** | 35-301 | 249693 | 149815800 | 120919 | 1263 | 0.167 |
| **BCHFD** | 35-301 | 340325 | 204195000 | 144057 | 1695 | 0.071 |
| **ITS 18S RNA Analyses** | | | | | | |
| **BND** | 35-301 | 243714 | 146228400 | 61774 | 22 | 0.501 |
| **BCND** | 35-301 | 205305 | 123183000 | 96481 | 28 | 0.457 |
| **BHFD** | 35-301 | 233519 | 140111400 | 90977 | 78 | 0.485 |
| **BCHFD** | 35-301 | 267408 | 160444800 | 74611 | 25 | 0.638 |

### Supplementary Table 3: Percent abundance of bacterial and fungal species in each samples

| Bacteria | Top 20 | Genus/Species | BND | BCND | BHFD | BCHFD |
|---|---|---|---|---|---|---|
| Genus | 1 | ***Lactobacillus*** | 0.208014566 | 59.74424552 | 64.94767575 | 15.75676749 |
| | 2 | ***Bacteroides*** | 0.25457188 | 18.0728148 | 12.54032302 | 2.06382536 |
| | 3 | ***Streptococcus*** | 0.016578556 | 0.38664059 | 0.637513806 | 20.01116093 |
| | 4 | ***Staphylococcus*** | 0.002044369 | 0.076726343 | 0.104976544 | 19.16652479 |
| | 5 | ***Pediococcus*** | 0.221175193 | 0.00752219 | 0.029524653 | 0.002837523 |
| | 6 | ***Desulfovibrio*** | 0.016211209 | 0.991424703 | 1.971590723 | 10.29926414 |
| | 7 | ***Clostridium*** | 0.004488029 | 4.054460659 | 0.115911601 | 8.299755973 |
| | 8 | ***Parabacteroides*** | 0.034738305 | 0.203099142 | 7.445680106 | 4.955261714 |
| | 9 | ***Enterococcus*** | 0.069412723 | 0 | 0.104976544 | 3.254639351 |
| | 10 | ***Escherichia*** | 1.59716E-05 | 5.077478562 | 3.993482706 | 0.001891682 |
| | 11 | ***Adlercreutzia*** | 0.061634537 | 1.712050549 | 1.751796083 | 3.268826968 |
| | 12 | ***Oscillospira*** | 0.025187267 | 0.544606589 | 1.054139466 | 6.063787526 |
| | 13 | ***Odoribacter*** | 0.00253949 | 2.480818414 | 1.595424772 | 1.029075157 |
| | 14 | ***AF12*** | 0.005590072 | 0.105310666 | 0.790604599 | 1.43105764 |
| | 15 | ***Weissella*** | 0.033764035 | 0 | 0.010935057 | 0.000945841 |
| | 16 | ***Helicobacter*** | 0.001309674 | 0 | 1.903793371 | 0.152280423 |
| | 17 | ***Flexispira*** | 0.003849164 | 1.499924778 | 0.062329823 | 0.472920568 |
| | 18 | ***Rothia*** | 0.000255546 | 0.398676094 | 0.215420617 | 1.117038382 |
| | 19 | ***Prevotella*** | 0.008512881 | 1.594704378 | 0.008748045 | 0.001891682 |
| | 20 | ***Akkermansia*** | 0 | 0 | 0 | 0.84085277 |
| Species | 1 | ***Bacteroides_acidifaciens*** | 37.04065041 | 8.487563362 | 53.05786003 | 3.785554936 |
| | 2 | ***Pediococcus_acidilactici*** | 49.18518519 | 0.035364847 | 0.138268454 | 0.020914668 |
| | 3 | ***Desulfovibrio_C21_c20*** | 0 | 6.294942827 | 1.063603489 | 64.82152817 |
| | 4 | ***Parabacteroides_distasonis*** | 2.09936766 | 0.141459389 | 24.95745586 | 22.7481874 |
| | 5 | ***Escherichia_coli*** | 0.003613369 | 39.78545326 | 19.4213997 | 0.013943112 |
| | 6 | ***Clostridium_butyricum*** | 0 | 30.81457032 | 0.015954052 | 0.013943112 |
| | 7 | ***Weissella_cibaria*** | 7.551942186 | 0 | 0.053180174 | 0 |
| | 8 | ***Akkermansia_muciniphila*** | 0 | 0 | 0 | 6.19771333 |
| | 9 | ***Ruminococcus_gnavus*** | 2.196928636 | 13.16751149 | 0.388215273 | 0.969046291 |
| | 10 | ***Mucispirillum_schaedleri*** | 1.416440831 | 0.04715313 | 0.186130611 | 0 |
| | 11 | ***Alistipes_indistinctus*** | 0.256549232 | 0.011788282 | 0.116996384 | 0.104573341 |
| | 12 | ***Butyricicoccus_pullicaecorum*** | 0.021680217 | 0.023576565 | 0.074452244 | 0.543781372 |
| | 13 | ***Streptococcus_infantis*** | 0.032520325 | 0.070729695 | 0.106360349 | 0.139431121 |
| | 14 | ***Serratia_marcescens*** | 0.104787715 | 0 | 0.069134227 | 0.006971556 |
| | 15 | ***Aggregatibacter_pneumotropica*** | 0 | 0.459743015 | 0 | 0.006971556 |
| | 16 | ***Propionibacterium_acnes*** | 0.014453478 | 0.212189084 | 0.005318017 | 0.104573341 |
| | 17 | ***Clostridium_methylpentosum*** | 0.007226739 | 0 | 0.159540523 | 0.125488009 |
| | 18 | ***Kocuria_palustris*** | 0.003613369 | 0.023576565 | 0.005318017 | 0.055772448 |
| | 19 | ***Jeotgalicoccus_psychrophilus*** | 0 | 0 | 0 | 0.069715561 |
| | 20 | **Acinetobacter_rhizosphaerae** | 0 | 0 | 0 | 0.062744004 |

| Fungi | Top 20 | Genus/Species | BND | BCND | BHFD | BCHFD |
|---|---|---|---|---|---|---|
| Genus | 1 | ***Aspergillus*** | 0 | 0.017663989 | 0.620482598 | 22.68181438 |
| | 2 | ***Candida*** | 2.168346676 | 99.59165013 | 91.88702324 | 12.12444556 |
| | 3 | ***Cystobasidium*** | 0.07130587 | 0 | 0.062270655 | 0 |
| | 4 | ***Debaryomyces*** | 0 | 0.010390582 | 0 | 0.046028956 |
| | 5 | ***Fusarium*** | 0 | 0.003117175 | 0.103413766 | 0.380785003 |
| | 6 | ***Issatchenkia*** | 0 | 0 | 0.064494607 | 0 |
| | 7 | ***Leptobacillium*** | 0 | 0 | 0.064494607 | 0 |
| | 8 | ***Malassezia*** | 0 | 0.008312466 | 0.045591015 | 0 |
| | 9 | ***Meyerozyma*** | 0.113441156 | 0 | 0.02223952 | 0 |
| | 10 | ***Millerozyma*** | 0 | 0 | 0.041143111 | 0 |
| | 11 | ***Penicillium*** | 0 | 0 | 0.114533526 | 0 |
| | 12 | ***Rhizopus*** | 0 | 0 | 0 | 1.72399364 |
| | 13 | ***Rhodotorula*** | 65.44096198 | 0 | 3.959746469 | 0 |
| | 14 | ***Saccharomyces*** | 0 | 0 | 0.186811965 | 0 |
| | 15 | ***Schizophyllum*** | 0 | 0.116374518 | 0 | 0 |
| | 16 | ***Syncephalastrum*** | 0 | 0 | 0 | 1.437358775 |
| | 17 | ***Talaromyces*** | 0 | 0 | 0.037807183 | 0 |
| | 18 | ***Trichosporon*** | 32.13787962 | 0.161054021 | 1.227621483 | 61.56791363 |
| | 19 | ***Wallemia*** | 0 | 0.020781164 | 0.471477816 | 0 |
| | 20 | ***Xeromyces*** | 0 | 0.058187259 | 0.879573001 | 0 |
| Species | 1 | ***Aspergillus_ficuum*** | 0 | 0 | 0 | 0.226299694 |
| | 2 | ***Aspergillus_ruber*** | 0 | 0 | 0.133450951 | 45.12538226 |
| | 3 | ***Candida_albicans*** | 2.284339607 | 99.78345965 | 93.41679672 | 35.37003058 |
| | 4 | ***Candida_hyderabadensis*** | 0.564165886 | 0 | 0 | 0.073394495 |
| | 5 | ***Cystobasidium_minuta*** | 0.061738908 | 0 | 0.018095044 | 0 |
| | 6 | ***Hyphopichia_burtonii*** | 0.014902495 | 0 | 0 | 0.048929664 |
| | 7 | ***Issatchenkia_orientalis*** | 0 | 0 | 0.065594535 | 0 |
| | 8 | ***Leptobacillium_leptobactrum*** | 0 | 0 | 0.065594535 | 0 |
| | 9 | ***Malassezia_restricta*** | 0 | 0.008328475 | 0.046368551 | 0 |
| | 10 | ***Meyerozyma_caribbica*** | 0.149024951 | 0 | 0.022618805 | 0 |
| | 11 | ***Millerozyma_farinosa*** | 0 | 0 | 0.04184479 | 0 |
| | 12 | ***Mucor_plumbeus*** | 0 | 0 | 0 | 0.06116208 |
| | 13 | ***Penicillium_hetheringtonii*** | 0 | 0 | 0.113094026 | 0 |
| | 14 | ***Rhizopus_arrhizus*** | 0 | 0 | 0 | 5.039755352 |
| | 15 | ***Rhodotorula_mucilaginosa*** | 85.9682364 | 0 | 4.027278279 | 0 |
| | 16 | ***Schizophyllum_commune*** | 0 | 0.116598651 | 0 | 0 |
| | 17 | ***Syncephalastrum_monosporum*** | 0 | 0 | 0 | 0.159021407 |
| | 18 | ***Trichosporon_insectorum*** | 10.85540322 | 0 | 0.390174391 | 13.89602446 |
| | 19 | ***Wallemia_tropicalis*** | 0 | 0.020821188 | 0.479518672 | 0 |
| | 20 | ***Xeromyces_bisporus*** | 0 | 0.058299325 | 0.894573749 | 0 |
